# Phenotypic heterogeneity in particle size is a viral mechanism of persistence

**DOI:** 10.1101/843177

**Authors:** Tian Li, Zhenyu Li, Erin E. Deans, Eva Mittler, Meisui Liu, Kartik Chandran, Tijana Ivanovic

## Abstract

Assembly of many enveloped animal viruses yields a mixture of particle morphologies, from small, essentially isometric forms to vastly longer, filamentous forms. Selective advantage of pleomorphic virus structure is apparent only *in vivo*, hindering functional characterization of distinct particle shapes. Here we sought to mimic the *in vivo* pressures on virus entry in cultured cells and in single-particle experiments of membrane fusion for influenza virus preparations enriched in spherical or filamentous particles. We show that filamentous shape confers functional advantage in the presence of neutralizing antibodies or fusion inhibitors and in cases of only limited fusion-protein activation. For very long particles, inactivation of >95% of associated fusion proteins still permits enough active-protein cooperation to induce membrane merger. Experiments with Ebola virus-like particles show that resistance to antibody pressure is a conserved feature of filamentous particles. Our results offer a strategy for averting drug resistance or immune evasion by targeting filamentous virus particles.

## INTRODUCTION

In 1946 Mosley and Wyckoff reported that influenza virus displays filamentous in addition to spherical particles (Mosley and Wyckoff, 1946). In the decades that followed, filamentous particles were reported for a number of circulating viruses including Newcastle disease virus, respiratory syncytial virus, and measles virus (Kilham et al., 1951; Liljeroos et al., 2013; Nakai and Imagawa, 1969). Some of the longest filamentous particles, which reached 500 µm in length, were reported for influenza C virus (Nishimura et al., 1990). Filamentous particle shape also characterizes a number of emerging pathogens, such as highly pathogenic avian influenza (HPAI), Ebola, Nipah and Hendra viruses (Bharat et al., 2012; Flewett and Challice, 1951; Hyatt et al., 2001; Liljeroos et al., 2013). While structural determinants of the filamentous-particle assembly have been defined for many viruses (Bialas et al., 2014; Bourmakina and Garcia-Sastre, 2003; Meshram et al., 2016; Noda et al., 2002; Roberts et al., 1998), function of the filamentous particle shape remains unknown.

The most basic function of virus particles is to spread their infectious cargo within and between hosts, and to initiate infection of new cells. Most published work attempting to define the function of virus particle shape in infection by pleomorphic viruses has focused on influenza. In early work, Chu et al. showed that the filamentous morphology is a phenotypic trait, and infection of chick embryos by influenza virus filtrates devoid of filamentous particles still yielded filamentous forms (Chu et al., 1949). Recent evidence confirms this view and shows that a single cell infected by a single initiating influenza virus produces particles with the size distribution characteristic of a bulk infection (Vahey and Fletcher, 2019). While functional analysis of particle shape has been challenging due to its phenotypic nature, one hypothesis is that the production of different particle forms ensures that infection can be initiated under a range of external pressures. However, filamentous particle shape does not confer a measurable infectivity advantage to influenza virus under normal cell-culture conditions (Dadonaite et al., 2016; Roberts et al., 1998), and filamentous particles are replaced by spherical ones after several passages *in vitro* (Choppin et al., 1960). On the other hand, influenza filaments are advantageous and are selected for *in vivo* (Seladi-Schulman et al., 2013). To develop a hypothesis for the function of the filamentous particle shape, we examined the unique pressures that are encountered by the extracellular form of the virus and that limit viral spread *in vivo*.

An obvious pressure that exists only *in vivo* are antibodies that are generated against surface components of the virus particle as part of the immune response to viral infection. The protection afforded by antibodies is evident in the success of sera from convalescent patients in treating acute infections in the 2013 West African EBOV epidemic, and anecdotal evidence that it is effective in treating emerging HPAI infections (Kong and Zhou, 2006; Sahr et al., 2017; Wu et al., 2015; Zhou et al., 2007). Viral glycoproteins, such as influenza virus hemagglutinin (HA) and EBOV glycoprotein (GP) are displayed prominently on the viral particle surface and are the main focus of the antibody response. HA and GP provide specificity for cellular receptors and mediate fusion with the endosomal membrane, thus enabling delivery of the viral infectious cargo to the cytoplasm of the host cell. Antibodies binding to HA and GP neutralize infection by interfering with these functions during viral entry. One question then is whether pleomorphic shape influences the sensitivity of viruses to neutralization by antibodies.

A second set of *in vivo* pressures arises from the requirement for activation of the viral glycoprotein by proteolytic cleavage and subsequent encounter of an appropriate fusion trigger. For influenza virus, cleavage of the inactive HA precursor, HA0, into two disulfide-linked subunits, HA1 and HA2, is required to activate HA for endosomal membrane fusion. Cleaved/active HA is metastable; low pH in endosomes reduces the barrier for transition to the stable post-fusion conformation and serves as the fusion trigger (Bullough et al., 1994). The HA of seasonal influenza is cleaved by specialized proteases expressed in the respiratory tract (e.g. human airway trypsin-like protease, transmembrane protease, serine 2) (Bottcher et al., 2006). In contrast, the HA of HPAI viruses contains a multi-basic cleavage site (MBCS) that can be cleaved by ubiquitous, furin-like serine proteases (Horimoto et al., 1994; Stieneke-Grober et al., 1992). Ubiquitous HA cleavage leads to more widespread activation of HPAI HA and contributes to systemic spread of HPAI viruses (Schrauwen et al., 2012). This phenomenon is uniquely observed *in vivo* because the use of trypsin for *in vitro* culture of cells results in complete HA cleavage.

Unlike for influenza virus, filamentous filovirus particles are more infectious than the spherical particles *in vitro*, demonstrating the contribution of the filovirus particle shape to infectivity even in the absence of the immune pressure (Welsch et al., 2010). EBOV, a filovirus, has a complex scheme for GP activation and triggering that includes extensive trimming by endosomal proteases and subsequent binding to the endosomal receptor, NPC1 (Carette et al., 2011; Chandran et al., 2005; Miller et al., 2012; Spence et al., 2016). A GP mutation that relaxes the host-factor dependency of membrane fusion conferred a selective advantage to EBOV during the 2013 West African epidemic (Wang et al., 2017). Pressures on the viral membrane fusion machinery might thus limit *in vivo* infections by diverse viruses, and might be greater for filoviruses because of their elaborate activation/triggering scheme than for influenza virus that senses a ubiquitous low-pH trigger. A second question, then, is whether pleomorphic virus particle shape affects the requirement for activation of the fusion glycoprotein.

In this study, we sought to examine how the pressures on viral entry posed by neutralizing antibodies and fusion glycoprotein activation differ in their effects on spherical and filamentous particles. We show that the filamentous particle shape confers an infectivity advantage in the presence of neutralizing antibodies, fusion inhibitors, and at very low levels of fusion-protein activation. A combination of single-particle membrane fusion experiments and stochastic simulations revealed that tens-of-micrometers long virus particles are fully resistant to inhibition at the level of membrane fusion. We furthermore demonstrate that the same basic principles apply to EBOV-like particles. We conclude that pleomorphic virus particles represent a general strategy enabling viral adaptation and persistence under changing environmental pressure on the viral cell-entry machinery.

## RESULTS

### Filamentous influenza virus particles have an infectivity advantage under HA pressure

We used filamentous influenza virus X31HA/Udorn (Ivanovic et al., 2013) and devised a strategy for particle size-enrichment using cycles of centrifugation at low relative centrifugal force, which preferentially sediments filamentous particles. We analyzed the resultant fractions by negative-stain electron microscopy (Figure 1A). The unfractionated X31HA/Udorn particles range in length from ∼50 nm to ∼2 µm (Figure 1B). Taking 250 nm as the cut off for filamentous particles, ∼17% of the unfractionated particles are filamentous. The supernatant fraction (SUP) has a relatively narrow and symmetric particle-size distribution, with 95% of the population shorter than 0.25 µm and 99.9% shorter that 0.75 µm. The pellet fraction (PELLET) is enriched in longer particles and includes 47.2% filaments. 13.1% of PELLET consists of particles longer than 0.75 µm (hereby designated as long filaments). The average length of filamentous particles in the PELLET is ∼5-times the average length of the combined SUP particles (Figure 1B, and see insets).

**Figure 1.**
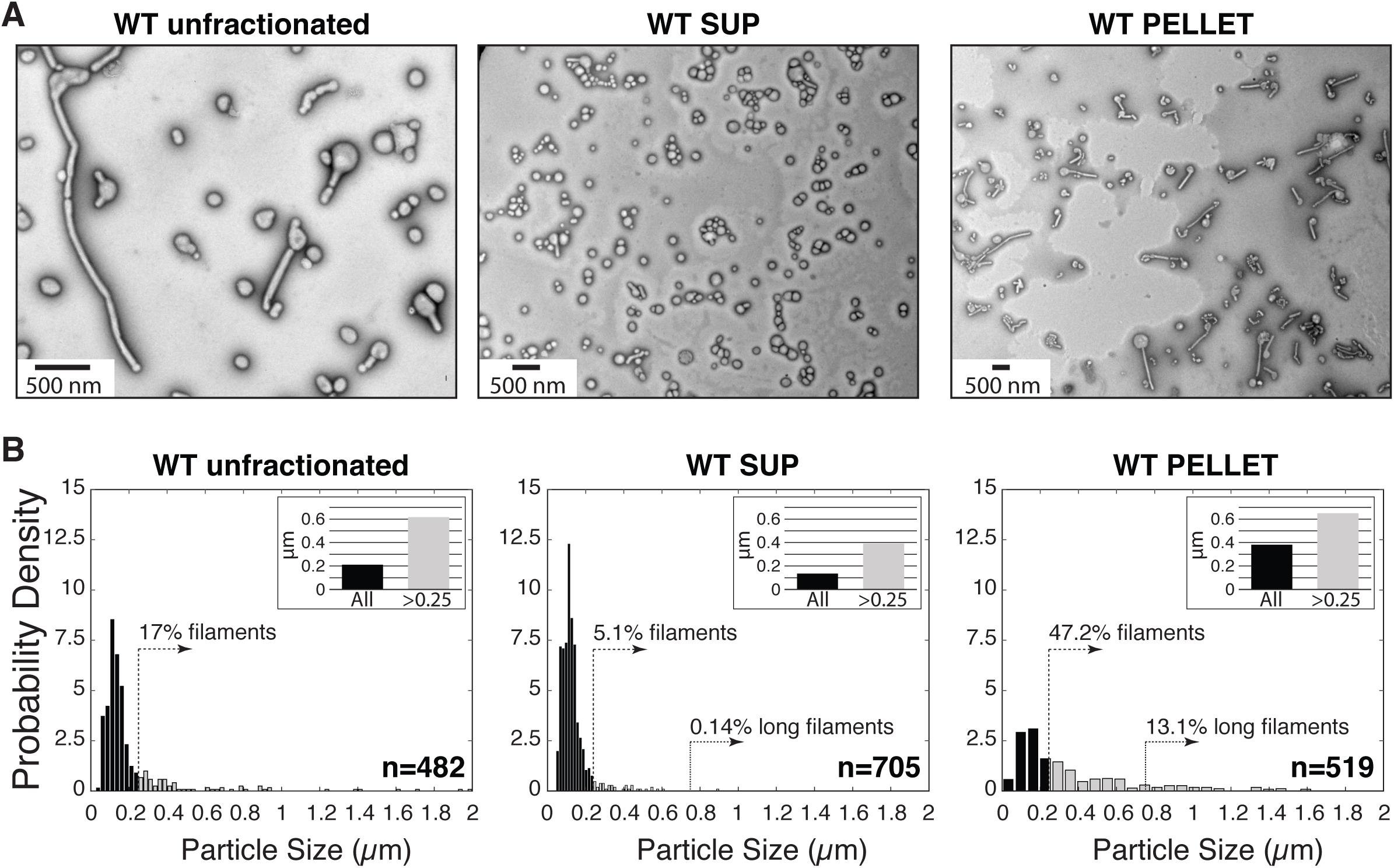
Particle fractions. (A) Sample negative-stain electron micrographs of WT X31HA/Udorn particles before and after fractionation into PELLET and SUP fractions. (B) Particle-length distributions derived from electron micrographs as those shown in (A). Length data were normalized so that the total area under the bars is equal to 1 on all plots. Dotted vertical lines mark 0.25 (filaments) or 0.75 µm (long filaments) length cutoffs, and the percentage of particles longer than the marked length is indicated above. Each inset shows mean particle length for the entire population (black bar), or the subset of the population longer than 0.25 µm (gray bar).

To determine whether viral filaments have an infectivity advantage under HA-directed selective pressures, we used (1) an HA-targeting broadly neutralizing antibody (IgG), HC19, which prevents HA-receptor binding (Bizebard et al., 1995), or (2) a Fab fragment of a broadly neutralizing antibody, MEDI8852 (M-Fab), which binds the base of HA and interferes with fusion-associated HA conformational changes (Kallewaard et al., 2016). We measured the SUP- and PELLET-particle concentrations (in hemagglutination units per ml; HAU/ml) needed to induce 50% cell death either in the presence of HC19 IgG or M-Fab in a single-cycle of infection (scTCID_50_) (Figure 2). We ensured that the HAU measurements accurately report on the relative particle counts for the PELLET and SUP fractions by quantitative EM (see Materials and Methods). We limited infections to a single cycle because particle-size enrichment is possible only for the initial infection, which then yields pleomorphic progeny (Chu et al., 1949; Vahey and Fletcher, 2019). In the absence of inhibition, the PELLET has no infectivity advantage and in some experiments a subtle disadvantage over the SUP (Figure 2C), consistent with previous reports for viral filaments (Dadonaite et al., 2016; Roberts et al., 1998). However, the effect of either HC19 IgG or M-Fab on the scTCID_50_ of the PELLET is lower than its corresponding effect on the SUP. The advantage of the PELLET over the SUP in the presence of HC19 IgG can at least in part be explained by the greater attachment of PELLET particles to cell-surface receptors (Figure S1A). The PELLET advantage in the presence of M-Fab ranges from 50% (17 nM M-Fab) to greater than 3-fold for the highest M-Fab concentration tested (268 nM) (Figure 2C and D). M-Fab does not affect attachment of particles to cells, so the PELLET also resists inhibition at the level of membrane fusion. Since short influenza particles are internalized via clathrin-mediated endocytosis (Rossman et al., 2012) whereas long filamentous particles are internalized by micropinocytosis, we also examined the effects of M-Fab on viral entry in the presence of the macropinocytosis inhibitor EIPA. Inhibition of micropinocytosis reduced the PELLET advantage in the presence of M-Fab (Figure S1B), confirming that the advantageous phenotype of the PELLET derives from cell entry by the filamentous particles in the mix. Taken together, these data confirm that virus particle shape modulates the sensitivity of influenza virus to HA inhibition both at the level of particle attachment to target cells and at the level of membrane fusion in endosomes.

**Figure 2.**
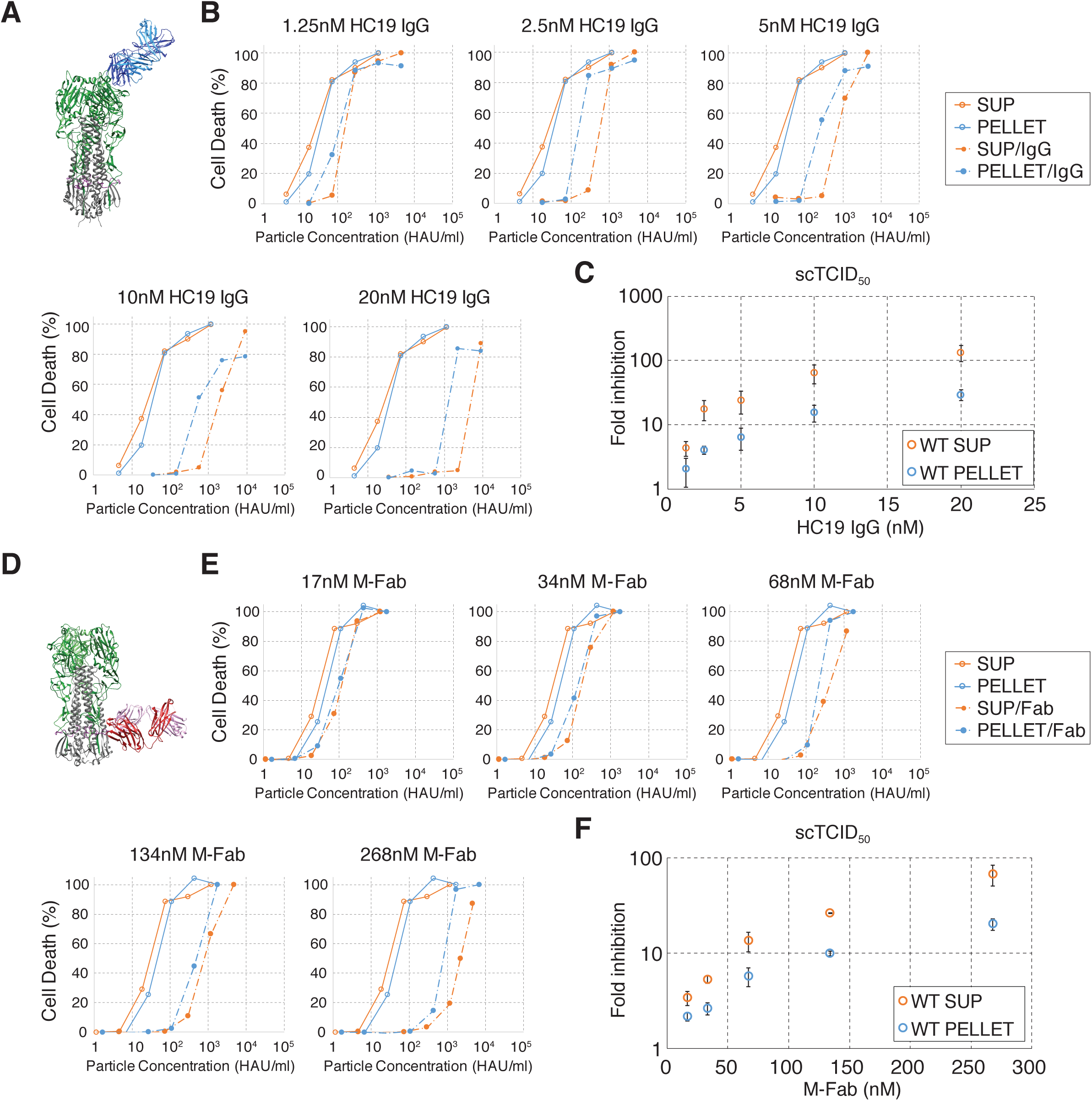
Filamentous virus particles have infectivity advantage when HA functions in receptor attachment or membrane fusion are inhibited. (A) Ribbon representation of X31-HA trimer in complex with the Fab fragment of the HC19 antibody (Bizebard et al., 1995). HA1 is shown in green, HA2 in grey, fusion peptide in magenta, and heavy and light chains of HC19 Fab in shades of blue. The trimeric HA image was assembled by overlaying the structure of the HC19 Fab in complex with an X31-HA1 monomer (PDB: 2VIR) with a structure of X31-HA trimer from a different source (PDB: 1EO8). (B) A representative set of scTCID_50_ measurements at various HC19 IgG concentrations. The no-inhibitor baseline for the PELLET and the SUP from which infectivity shifts are measured in (C) is shown on all plots. (C) Fold increase in scTCID_50_ for samples treated with HC19 IgG relative to untreated samples. Average and standard deviation of three independent experiments are shown. (D) Ribbon representation of an H7-HA trimer in complex with M-Fab (PDB: 5JW3) (Kallewaard et al., 2016). HA1 is shown in green, HA2 in grey, fusion peptide in magenta, and heavy and light chains of M-Fab Fab in red and pink. (E) A representative set of scTCID_50_ measurements at various M-Fab concentrations. The no-inhibitor baseline for the PELLET and the SUP from which infectivity shifts are measured in (F) is shown on all plots. (F) Fold increase in scTCID_50_ for samples treated with M-Fab relative to untreated samples. Average and standard deviation of three independent experiments are shown.

### Filamentous influenza virus particles fuse more efficiently when HA activity becomes limited by presenting more opportunities for HA insertion

To investigate potential mechanisms underlying the effects of viral particle shape on membrane fusion, we took advantage of prior work by our group and others characterizing HA-mediated membrane fusion of influenza virus. This process results from a functional interplay among HAs in the contact patch between the particle and the target membrane. There are ∼50 HAs in the contact patch for the smallest spherical particles, and this number scales linearly with particle length (Calder et al., 2010; Ivanovic et al., 2013; Wasilewski et al., 2012). Low pH triggers a conformational change in HA leading to insertion of its fusion peptides (FP) into the target membrane (productive HA). If HA assumes the post-fusion state without engaging the target membrane, FP inserts back into the viral membrane inactivating HA (unproductive HA) (Ivanovic and Harrison, 2015; Wharton et al., 1995). HA triggering by low pH is stochastic, but the combined free energy released by conformational changes of 3-5 neighboring HAs (fusion cluster) is required to induce merging of the two membranes (Ivanovic et al., 2013; Ivanovic and Harrison, 2015). The unproductive as well as the uncleaved/inactive HAs impose restrictions on the available geometries for adjacent HA insertions influencing the rate and probability of fusion-cluster formation (Ivanovic and Harrison, 2015; Otterstrom et al., 2014). We hypothesized that the larger contact patch on filamentous virus particles contributes to their lower sensitivity to fusion inhibitors by presenting more opportunities for HA insertion.

To investigate the mechanism responsible at the level of membrane fusion for the resistance of filamentous particles to inhibition by M-Fab, we measured the kinetics of membrane fusion for the SUP and PELLET fractions of X31HA/Udorn in single-particle experiments (Figure 3A). Analysis of individual particles can reveal subpopulations with distinct kinetics, and in this way, obviate the need for homogeneous particle preparations. We performed membrane fusion experiments with DiD-labeled SUP or PELLET particles on supported planar membrane bilayers in the presence of a range of M-Fab concentrations at pH5.2 (Movie S1-S4). We extracted individual-particle lag times from the time of pH drop to the time of hemifusion, a fusion intermediate that reports on fusion-cluster formation and is detected by the onset of DiD dequenching (Figure 3A) (Floyd et al., 2008; Ivanovic et al., 2013). We fitted frequency distributions of hemifusion lag times with a gamma probability density and derived mean lag times from the fits (Figure 3B and C) (Floyd et al., 2008; Ivanovic and Harrison, 2015). The plots of hemifusion yield and time for a range of M-Fab concentrations both show that the PELLET particles are less sensitive to HA inactivation than the SUP particles (Figure 3C). We derived hemifusion yield and lag-time distributions for the long (>250 nm) particles within the PELLET (Figure 3D) as described in the Materials and Methods. In the absence of HA pressure, fusion efficiency is not limited by particle size. The advantage of filaments becomes apparent in the presence of M-Fab, then becomes more pronounced at higher M-Fab concentrations (Figure 3D, left). On the other hand, the difference in fusion rate between the short and long particles becomes smaller for higher M-Fab concentrations until the plateau value is reached (∼450 sec in our experiments) (Figure 3D, right). The combined results thus show that viral filaments fuse more rapidly at low HA pressures, and more efficiently at high HA pressures than short particles. To aid in interpreting the difference in sensitivity of long and short particles to M-Fab, we quantified binding of fluorescently tagged M-Fab (M-Fab^fl^) to unlabeled virus particles. M-Fab^fl^ bound to SUP particles exhibits a narrow distribution of intensities, while M-Fab^fl^ bound to PELLET particles has a wider fluorescence-intensity distribution with a long tail (Figure S2A). M-Fab binding thus scales with particle length (see Figure 1), suggesting that long particles escape M-Fab effects rather than M-Fab binding. Incubation of M-Fab^fl^-bound particles at low pH did not significantly change fluorescence-intensity distributions (Figure S2A, bottom), showing that M-Fab remains particle-bound for the duration of fusion experiments, and that it therefore eliminates targeted HAs from participating in the fusion reaction. Long particles thus tolerate levels of HA inactivation that inhibit fusion of shorter particles.

**Figure 3.**
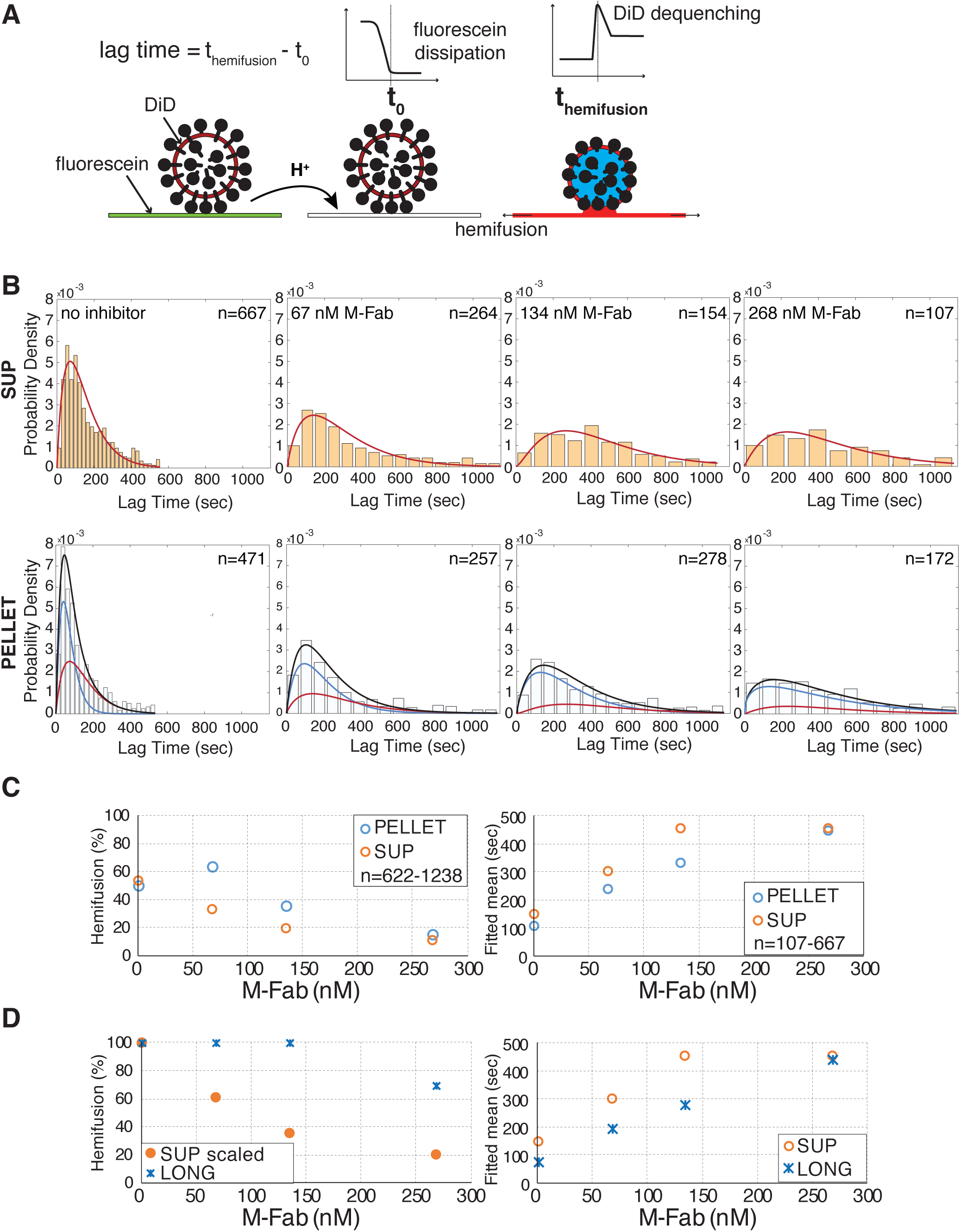
Filamentous particles fuse more rapidly and/or more efficiently depending on the extent of HA inactivation. (A) Schematic of the single-particle assay of membrane fusion (Floyd et al., 2008; Ivanovic et al., 2013). Particles attach to planar membrane bilayers by engaging sialic-acid receptors displayed on GD1a molecules in the membrane. Fluorescein is bound to the target membrane and serves as a pH sensor. DiD is incorporated in the viral membrane so that it is partially quenched. Hundreds of individual virus particles can be monitored in a single experiment using total internal reflection fluorescence microscope. Times from pH drop (dissipation of fluorescein fluorescence) in the entire field of view to hemifusion (onset of DiD dequenching) for individual detected particles are extracted from time-lapse movies and plotted as probability-density histograms (see panel (B)). (B) Hemifusion lag times with gamma-distribution fit (SUP, *red line*), or bi-gamma fit (PELLET, *black line*) with the underlying distributions for the short (*red line*) and long (*blue line*) PELLET-particle subpopulations. Data were pooled from several experiments where each condition was represented on each day. The total number of particles represented by each histogram is indicated on the plots. (C) Hemifusion yield, and fitted mean from panel (B). The total number of particles analyzed per each condition (a single data point) is indicated as the ‘n-value’ range. (D) Derived hemifusion yield and fitted mean for long particles in the PELLET (red and blue lines in panel (B), and see Materials and Methods).

A simple interpretation of the greater tolerance of long particles to HA inactivation is that the larger contact patch afforded by these particles increases the total number of inserted HAs for a given extent of HA inactivation. At low HA pressure, a larger number of inserted HAs would translate to a greater number of possible fusion clusters forming, and thus a shorter time until the first fusion cluster can form and induce hemifusion. At high HA pressure, a larger number of inserted HAs would translate to an increased likelihood of fusion clusters forming at all, and thus higher fusion efficiency (Figure 4A). To express these predictions in quantitative terms, we performed stochastic simulations of fusion using the algorithm we previously described (Ivanovic et al., 2013; Ivanovic and Harrison, 2015) for a 600-fold range of patch sizes and zero to 80-97% HA inactivation (Figure 4A and B). The chosen patches represent particles ranging from ∼50 nm in diameter (PS 55, or 55 HAs interfacing the target membrane) to ∼30 µm in length (PS 34627) (Ivanovic et al., 2013). The resultant trends (Figure 4B) are in close agreement with our single-particle measurements of membrane fusion in the presence of M-Fab (Figure 3D). At high HA activity levels, all patches have a high probability of assembling fusion clusters, but larger patches form fusion clusters more rapidly. The probability of fusion-cluster formation for the smallest patch declines for HA inactivation above 20% (Figure 4B, black line), but larger patches require greater HA inactivation before any reduction in the fusion-cluster yield. The rate differences for different sized patches show modest (<20%) increases with increasing levels of inactivation before converging at a plateau at higher levels of HA inactivation. Our measured trends of hemifusion time and yield for SUP versus long particles in the presence of M-Fab (Figure 3D) most closely match the simulation results for patch-size pairs in which the larger patch is ∼4-5 times the size of the smaller one (e.g. 583/121, or blue and orange lines in Figure 4B). In close agreement with the simulation, we obtained the value of 4.9 for the ratio between the average particle lengths for the long particles (>250 nm) in the PELLET and combined SUP particles (Figure 1, insets). The simulation results thus support the notion that filamentous virus particles fuse more rapidly and/or more efficiently in the presence of fusion inhibitors than do shorter particles, by having more opportunities for HA insertion and fusion-cluster formation.

**Figure 4.**
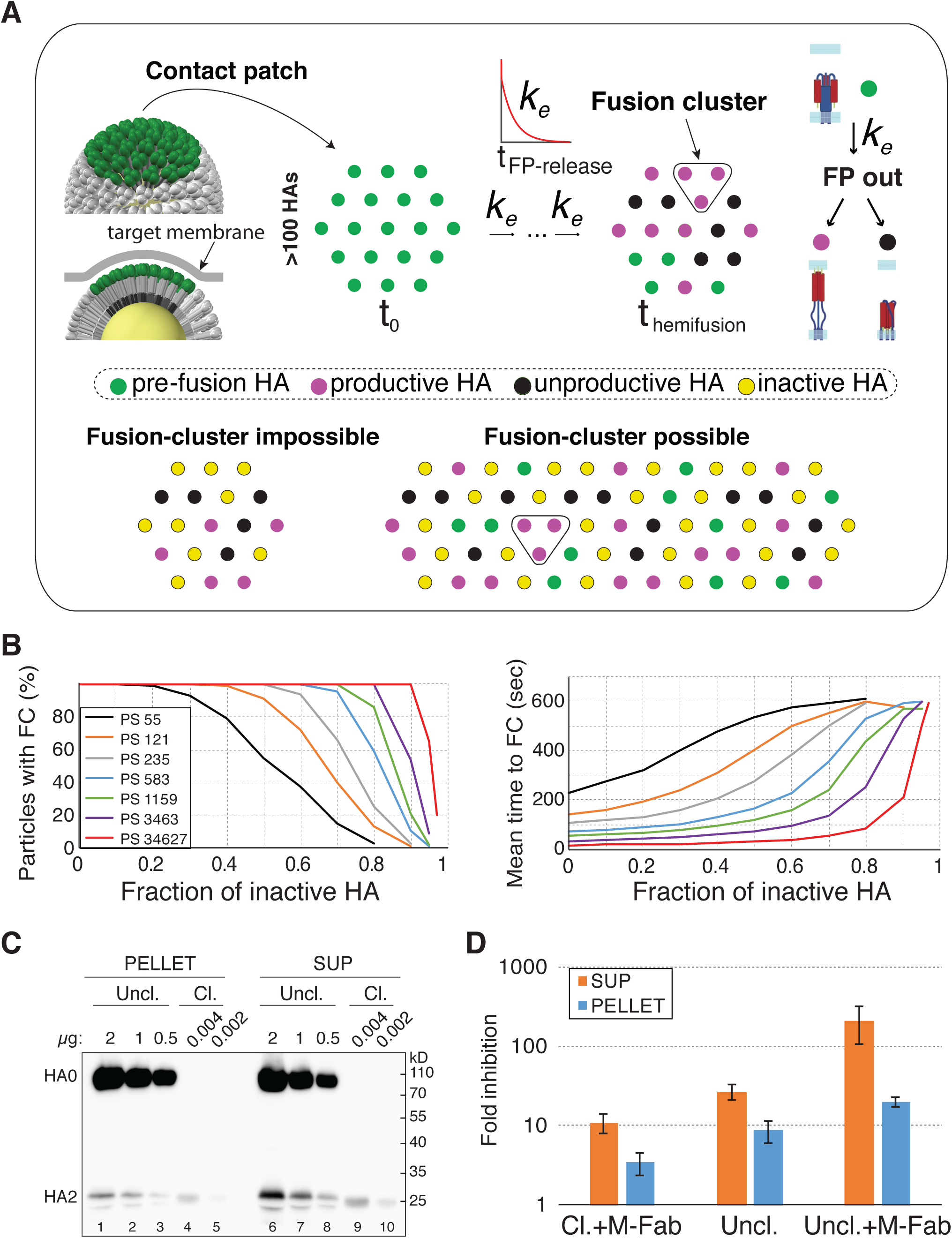
Filamentous particles are resistant to extreme HA inactivation. (A) Schematic of the stochastic fusion model (Ivanovic et al., 2013; Ivanovic and Harrison, 2015). *Top*: a portion of the virion surface contacting the target membrane, contact patch (green HAs on the displayed virion model shown as a top or a side view), is represented as a hexagonal lattice of HAs. A simplified contact patch before fusion (t_0_) or at the time of fusion-cluster formation (t_hemifusion_) is shown. Random HAs extend and insert into the target membrane (green – pre-fusion HA; magenta – inserted HA) or become inactivated (black – unproductive HA). *k*_*e*_ is the rate of FP release. *Bottom*: for the same frequency of unproductive and inactive HAs, longer particles are predicted to have a higher probability of fusion-cluster formation. Inactive (inhibitor bound or uncleaved) HAs are shown in yellow. The prediction for the advantage of filamentous particles is illustrated with an elliptical patch, but the simulations used circular patches (see Materials and Methods). (B) Simulation results for the yield and mean lag-time to fusion-cluster (FC) formation at 0.02-0.1 inactive HA-fraction increments from datasets with at least 1000 fusion clusters. PS, patch size represented with the number of HA molecules. (C) Western blot of purified virus particles. LC89 antibody detects an epitope in HA2 (Wharton et al., 1995). Total viral protein loaded in each lane is indicated (µg). To aid the estimate of the total HA0 cleavage in the uncleaved prep (Uncl.), a small amount of fully cleaved virus (Cl.) was loaded as a reference in lanes 4, 5, 9, and 10. (D) Fold reduction in infectivity (plaque-forming units) per input particle (HAU) for uncleaved and/or M-Fab treated samples, relative to cleaved, untreated samples. The experiment was set up in a way to ensure that any differences among samples result from the initial infection by the SUP and PELLET fractions (see Materials and Methods). Data is the average and standard deviation of three independent measurements.

### Long filamentous influenza virus particles resist extreme HA inactivation

The simulation results in Figure 4B show a striking prediction that the longest members in a pleomorphic virus population are effectively resistant to inhibition at the level of membrane fusion. A 30-µm long particle is predicted to require at least 90% HA inactivation for any reduction in the fusion yield, and retains 20% probability of fusion even with 97% inactive HAs (Figure 4B, red line). The probability of fusion for the shortest patch, on the other hand, goes to zero for HA inactivation levels greater than 80%. The simulation results and particle-size distributions of the PELLET and the SUP together predict that the infectivity advantage of the PELLET should be greater at higher levels of HA inactivation, as the probability of fusion goes to zero for shorter particles. This is because the SUP and PELLET fractions are more different for longer length cutoffs, and, for example, the PELLET has 10-times more filaments (> 0.25 µm) and 100-times more long filaments (> 0.75 µm) than the SUP (Figure 1B). Consistent with this prediction are the outcomes of the scTCID_50_ experiments, in which the PELLET advantage increased from 50% to 3-fold for increasing M-Fab concentration (Figure 2D).

To validate this experimentally, we sought to drive HA inactivation levels to the extreme by generating virus particles that retain nearly 100% uncleaved (inactive) HA0 (Figure 4C). Since uncleaved viruses are WT viruses produced in the absence of activating protease, we cannot exclude the possibility that some HA is cleaved during entry by endosomal proteases, but this effect is expected to be small at best. We measured the reduction of infectivity for the uncleaved virus, or the uncleaved virus pre-treated with 268 nM M-Fab, relative to fully cleaved, untreated virus. For this, we used a modified plaque-assay experiment to measure differences in infectivity during the first round of infection when infecting viruses differ in size. The result for the control (cleaved, M-Fab treated) virus matched the scTCID_50_ data at this M-Fab concentration (Figure 2C, D) with the PELLET offering approximately 3-fold higher infectivity than the SUP (Figure 4D). A similar result was obtained for the uncleaved virus suggesting that limited HA activation probably did occur in endosomes. Consistent with our prediction for extreme HA pressure, the uncleaved, M-Fab-treated PELLET is at least 10-times more infectious than the corresponding SUP. We estimate that the inactive HA fraction on the uncleaved, M-Fab-treated PELLET is greater than 95% (see Materials and Methods). Moreover, the PELLET infectivity is reduced only 20-fold at this extreme HA pressure, which we now can interpret to primarily result from the loss of infectivity of the shorter members of the population. Our results thus support the prediction that the longest subset of the population is resistant to extreme HA inactivation.

### Endosomal fusion by influenza virus particles is insensitive to fusion rate reduction resulting from HA inactivation

The release of FP from the pre-fusion pocket on HA is the rate-limiting structural rearrangement influencing the rate of fusion-cluster formation (Ivanovic et al., 2013). The fold-back for HAs in the fusion cluster is cooperative and fast (Ivanovic et al., 2013). Indistinguishable baseline infectivity of the SUP and the PELLET in the absence of HA pressure (Figure 2) suggests that infection does not benefit from the higher rate of fusion-cluster formation afforded by the long particles in the PELLET (Figure 3). However, since HA inactivation reduces rate differences between long and short particles (Figure 3 and 4), the preceding experiments do not address whether rate reduction resulting from HA inactivation contributes to infectivity outcomes.

To test directly whether changes in the kinetics of fusion-cluster formation influence infectivity, we used virus mutants with (de)stabilized pre-fusion HA. A mutation in HA2, D112_2_A, destabilizes FP in its pre-fusion pocket resulting in faster fusion-cluster formation (Ivanovic et al., 2013). A mutation in HA1, H17_1_Y, stabilizes FP in its pre-fusion pocket and is predicted to slow fusion-cluster formation (Thoennes et al., 2008). To aid the interpretation of infectivity experiments, we first compared the kinetics of membrane fusion for WT, D112_2_A, and H17_1_Y viruses in single-particle experiments. WT and D112_2_A gave very similar hemifusion-yield curves and a similar relative change in hemifusion lag-times with increasing M-Fab concentration (Figure 5A) owing to the same extent of M-Fab binding to mutant and WT particles (Figure S2B). However, D112_2_A particles fuse 7 to 11 times more rapidly for the tested range of M-Fab concentrations (Figure S3 and 5A) due to more rapid fusion-peptide release for the mutant (Ivanovic et al., 2013). At even the highest M-Fab concentration, hemifusion of D112_2_A particles is 2.4-fold faster than that of the uninhibited WT (Figure S3 and 5A). H17_1_Y, on the other hand, is so stabilized that it has lower hemifusion yield at pH 5.2 (Figure 5A). At pH 4.8, the hemifusion yield of H17_1_Y is comparable to that of WT and shows a similar trend with M-Fab (Figure 5B), owing to the comparable extent of M-Fab binding to the mutant and WT particles (Figure S2B). However, hemifusion is markedly slower for the mutant at all M-Fab concentrations, and even ∼2.2-fold slower in the absence of M-Fab than for the WT at 268 nM M-Fab (Figure S3 and 5B). Thus, the HA mutants allow us to probe the effects of altered hemifusion rates on infectivity independently of effects of M-Fab on fusion efficiency.

**Figure 5.**
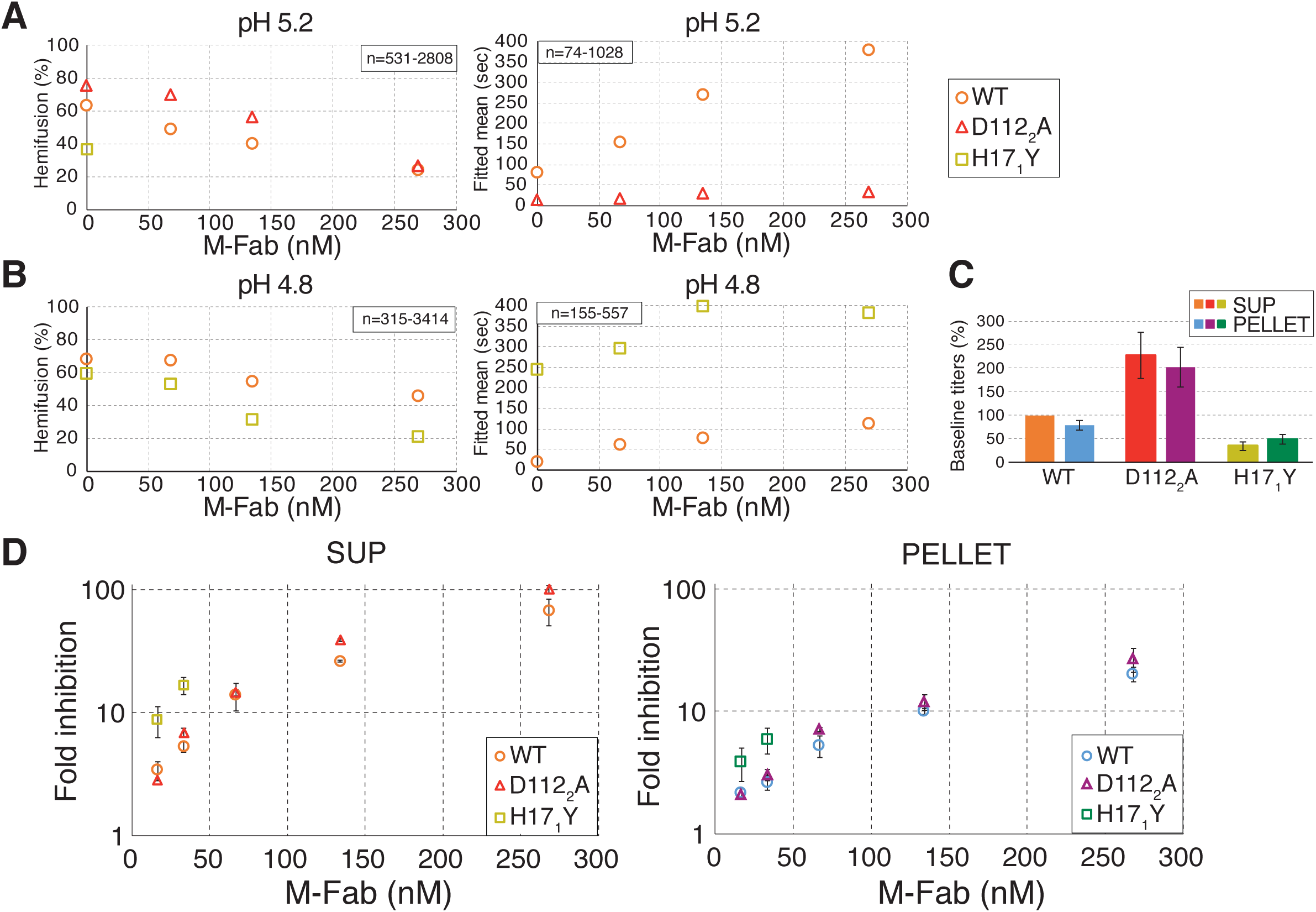
Endosomal fusion is insensitive to rate changes resulting from HA inhibition. (A and B) Single-particle hemifusion yield and fitted mean for a range of M-Fab concentrations for short particles at pH 5.2 (A) or pH 4.8 (B). The total number of particles analyzed per each condition (a single data point) is indicated as the ‘n-value’ range. (C) Baseline scTCID_50_ titers normalized to WT SUP. Average and standard deviation of three independent experiments are shown. (D and E) Fold increase in scTCID_50_ for M-Fab-treated samples relative to untreated samples for SUP (D) or PELLET (E) for WT, D112_2_A, and H17_1_Y viruses. Average and standard deviation of three independent experiments are shown. WT data is replotted from Figure 2F for direct comparison.

We next compared WT, D112_2_A, and H17_1_Y viruses (either PELLET or SUP) in scTCID_50_ experiments (Figure 5C, 5D, and S4). The baseline titers of D112_2_A and H17_1_Y viruses are within 2-3-fold of WT, showing that cell entry is relatively insensitive to the rate of fusion-cluster formation (Figure 5C). Furthermore, WT and D112_2_A viruses are equally sensitive to all tested M-Fab concentrations, showing that the rate increase afforded by HA destabilization does not boost infectivity under HA pressure (Figure 5D). H17_1_Y is more sensitive to HA inactivation than WT and D112_2_A viruses, but otherwise follows the same trend for the higher M-Fab concentration for both the PELLET and the SUP (Figure 5D). The higher sensitivity of H17_1_Y to M-Fab can be attributed to lower hemifusion yield of this mutant at the pH of endosomes (Figure 5A). If the lower rate of fusion-cluster formation for H17_1_Y contributed to its higher sensitivity to M-Fab, a further increase in M-Fab concentration would inhibit H17_1_Y disproportionately, causing it to deviate from the WT/D112_2_A infectivity curves. The same infectivity trend for H17_1_Y confirms that cell entry of even the slow-fusing H17_1_Y mutant is insensitive to further rate reduction caused by HA inactivation (Figure 5D). Consequently, the long particles in both the H17_1_Y and D112_2_A PELLET offer the same protection to HA inactivation as the long particles in the WT PELLET. HA inactivation thus lowers infectivity by reducing the number of particles that successfully assemble fusion clusters, and rate reduction is inconsequential for infectivity.

### Lower sensitivity of long particles to antibody pressure is a common property of pleomorphic viruses

To extend the observations for influenza virus to another pleomorphic viral pathogen, we generated Ebola virus-like particles (EBO VLPs) incorporating EBOV GP and the matrix protein VP40 fused at its N-terminus to *Renilla* luciferase. Unlike influenza virus which always packages a single genome independent of the particles size, filamentous EBOV particles can incorporate more than one genome (Beniac et al., 2012). Particles without genomes but enclosing a reporter protein thus allow us to focus any functional differences among particles of different sizes to cell entry. EBO VLPs formed authentic particle structures ranging from ∼70 nm to greater than 2 µm (Figure 6A). We performed similar cycles of low-force centrifugation as we did for influenza virus to enrich for short (EBO VLP SUP) and long (EBO VLP PELLET) particles. About 92% of the EBO VLP SUP particles are shorter than 250 nm, and about 58% of the EBO VLP PELLET particles are longer than 250 nm (Figure 6B). Taking the cut off of 250 nm, the average length of filamentous particles in the EBO VLP PELLET is 3.3-times the average length of the combined SUP particles (Figure 6B, and see insets).

**Figure 6.**
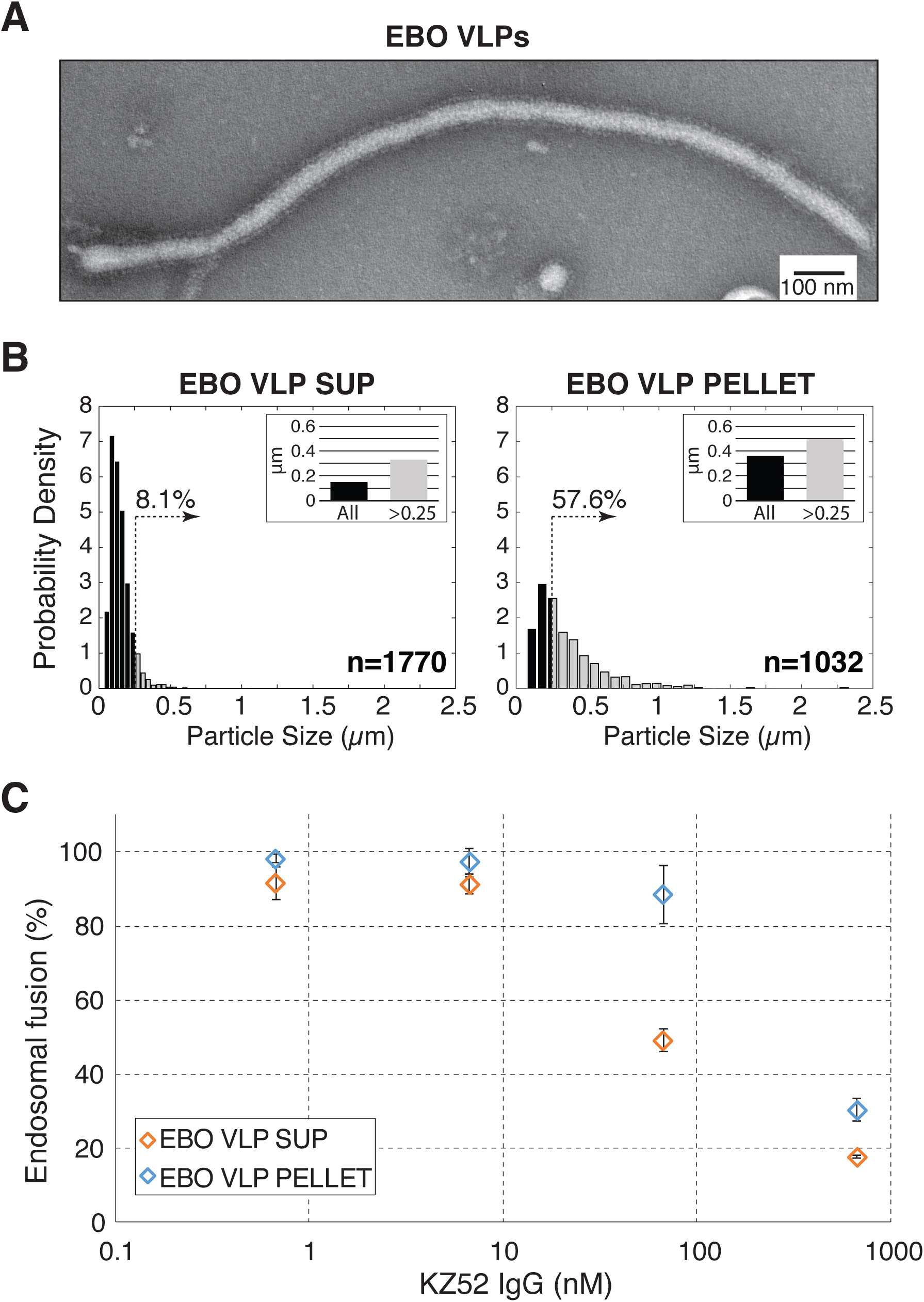
Lower sensitivity of filamentous particles to neutralizing antibodies is a general property of pleomorphic viruses. (A) A filamentous and a spherical EBO VLP, each showing surface decorated with EBOV GP. (B) Particle-length distributions for the EBO VLP SUP and PELLET derived from electron micrographs. Length data were normalized so that the total area under the bars is equal to 1. Dotted vertical lines mark the 0.25-µm length cutoff, and the percentage of particles longer than 0.25 µm is indicated above. Insets show mean particle length for the entire population (black bar), or the subset of the population longer than 0.25 µm (gray bar). (C) Cytoplasmic luciferase activity reports on endosomal membrane fusion by EBO VLPs incorporating luciferase-VP40 protein. KZ52 IgG is more effective against the SUP than the PELLET particles. Average and standard deviation of three independent experiments are shown.

We measured the extent of endosomal membrane fusion by entering EBO VLPs by quantifying cytoplasmic luciferase activity in live cells 6.5 hours post infection. Since EBO VLPs cannot proceed with steps in infection past the initial cell entry, measured effects are limited to the initial infection analogous to influenza virus scTCID_50_ experiments (Figure 2). We treated the VLPs with a neutralizing antibody KZ52, which binds the base of the EBOV GP, and clamps GP1 and GP2 subunits together (analogous to clamping the HA1 and HA2 subunits of HA) (Lee et al., 2008). KZ52 IgG does not affect virus particle attachment to cells or endosomal cleavage by cathepsins, thus its effects are solely focused on endosomal membrane fusion. EBO VLPs showed the expected sensitivity to KZ52 IgG, but the PELLET was less sensitive than the SUP. Long particles in the EBO VLP PELLET thus display greater tolerance to fusion inhibition analogous to our findings for influenza virus (Figure 2). Lower sensitivity of filamentous particles to antibody pressure is thus a general property of pleomorphic viruses.

## DISCUSSION

Our results for influenza virus and EBO VLPs reveal the role of filamentous particles in permitting cell entry under pressures on the viral entry machinery. We demonstrate the infectious advantage of filamentous virus particles in the presence of neutralizing antibodies (Figure 2A and 6C) or fusion inhibitors (Figure 2B and 6C) or at very low levels of fusion-protein activation (Figure 4D). Very long filamentous particles are resistant to extreme inactivation, and greater than 95% of the particle-associated viral glycoproteins can be rendered inactive without significant consequences on infectivity (Figure 4). Long viral filaments thus resist any pressure that individually targets components of the viral cell-entry machinery. Since heterogeneity in virus-particle size is a phenotypic property, filamentous virus particles enable a pathway of adaptation that is not initiated by genetic change.

Together, these findings lead us to a model in which filamentous particles enable established infections, such as those by influenza virus, to persist in circulation under changing environmental pressure (Figure 7). At low relative selective pressure, most new influenza virus infections are initiated by spherical particles (Figure 7A); they are more numerous, use fewer resources, and are faster to internalize and assemble (Dadonaite et al., 2016; Vahey and Fletcher, 2019). Filamentous particles take over when cell-entry is limited, enabling replication and ensuring subsequent rounds of infection (Figure 7B). If selective pressure is removed, the genetically identical viral population can expand. Under sustained pressure, viral replication initiated by filamentous particles can lead to (genetic) adaptation.

**Figure 7.**
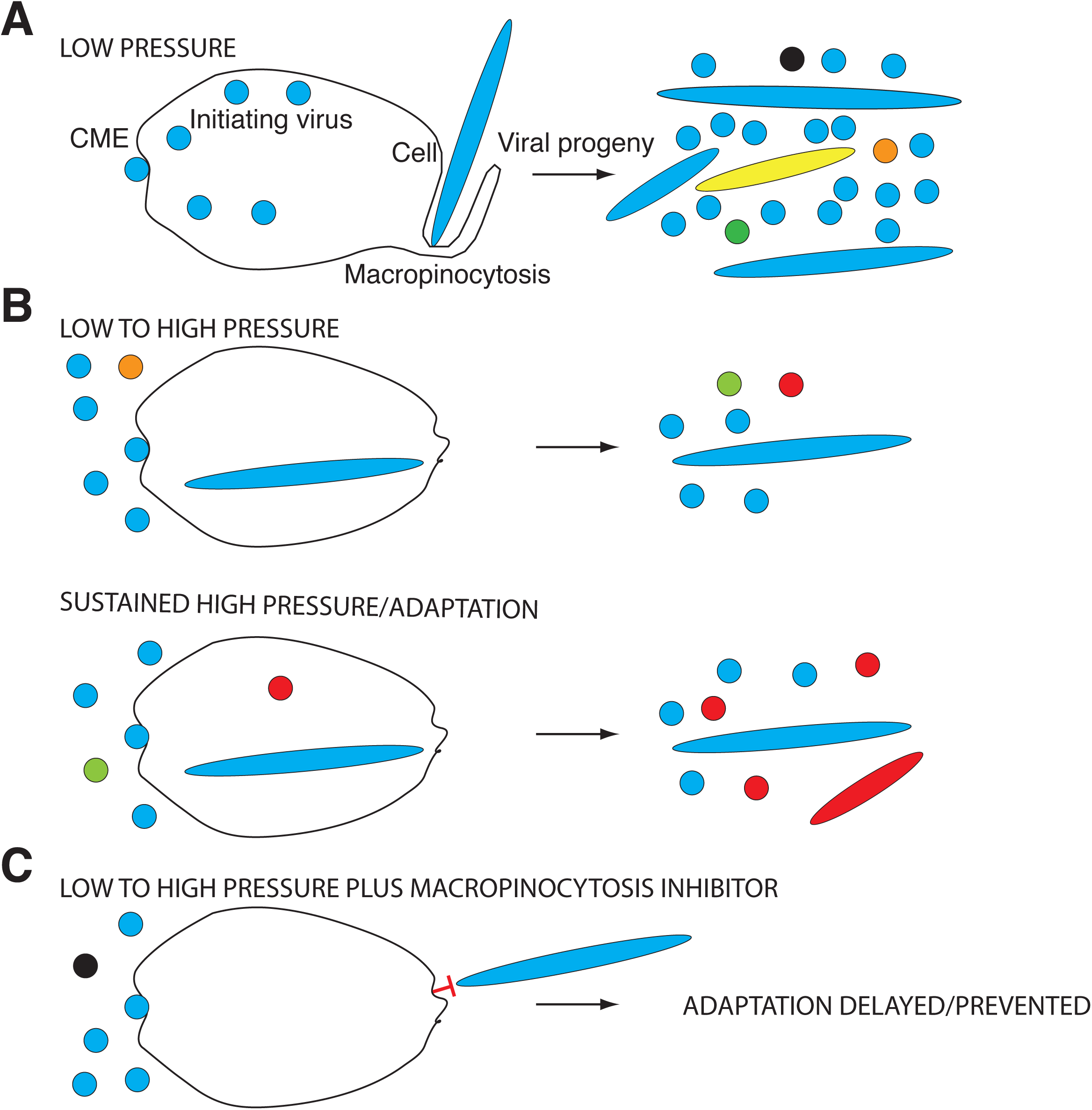
A model for the role of filamentous particles in viral persistence. Viral progeny is pleomorphic independent of the initiating-virus morphology. Random mutations arise during replication. (A) At low pressure on cell-entry machinery, infections by spherical particles dominate (circles, spherical particles; ellipses, filaments; blue, WT particles). Spherical particles might become internalized via clathrin-mediated endocytosis (CME) as in the case of influenza virus, or macropinocytosis as in the case of Ebola virus (Nanbo et al., 2010; Rossman et al., 2012). Filamentous particles invariably require macropinocytosis for internalization. Random mutations arise during virus replication (different color particles). (B) Filamentous particles can enter cells permitting some level of replication when spherical particles are unable, and can by random chance lead to adaptive mutations (red color particle). (C) Preventing filamentous particle assembly or cell entry (e.g. by using macropinocytosis inhibitors) might avert viral adaptation to immune pressure or man-made inhibitors. The same strategy might apply to preventing zoonotic adaptations that can lead to pandemics.

Our results also reveal a pathway of adaptation for emerging infections, such as those by Ebola virus (Figure 7B). A special case of selective pressure on cell entry machinery arises from zoonotic transmissions that are accompanied by a period of low cell-entry efficiency as viruses adapt to novel, host-specific conditions, e.g. different receptors and/or fusion-specific factors (Galloway et al., 2013; Gamblin and Skehel, 2010; Wang et al., 2017). In this context also, filamentous particles might offer a pathway for adaptation by compensating for functional deficiencies of cell-entry proteins in a new host and permitting their repair through subsequent mutation (Figure 7B). Since all filamentous particles use a common cellular pathway (macropinocytosis) for internalization into cells (de Vries et al., 2011; Krzyzaniak et al., 2013; Nanbo et al., 2010; Pernet et al., 2009; Rossman et al., 2012; Saeed et al., 2010), future dissection of host-factor requirements in this process could lead to the development of broad-spectrum antiviral treatments with potential to curb pandemics by diverse emerging pathogens such as HPAI, EBOV, Nipah, and Hendra viruses (Figure 7C).

Development of a next generation influenza virus vaccine has been a top priority in influenza vaccine research. An ideal vaccine would eliminate the need for annual revaccinations against seasonal influenza and also offer protection from potential pandemic strains. While diverse strategies are being explored for eliciting broad protection against influenza virus infection, all HA-targeting efforts focus on eliciting antibodies specific for conserved and functionally constrained epitopes involved either in receptor binding or membrane fusion. HA-targeting antibodies also offer protection by mechanisms not limited to inhibition of virus cell entry (Vogel and Manicassamy, 2020). For example, stem-binding antibodies primarily target HA on virus-infected cells and induce antibody-dependent cell-mediated cytotoxicity (ADCC) (DiLillo et al., 2014; He et al., 2016). Nonetheless, independent of the antibody mechanism of action, mutant HAs that escape recognition by the broadly neutralizing antibodies are expected to be deficient in cell-entry functions, because simultaneous acquisition of resistance and permissive mutations is a low-probability event in the absence of selective pressure for the permissive mutations. As we have shown here, if filamentous particles can compensate for functional deficiencies in HA, even the low level of replication initiated by filamentous particles harboring resistant mutations could ultimately lead to repair of function through secondary mutation. Filamentous particles thus represent a potential weakness of HA-targeting treatments or vaccination strategies by offering a pathway for establishing a functional epistasis that rescues deleterious effects of resistance mutations.

Our results offer a new paradigm for achieving lasting inhibition of pleomorphic virus infection by targeting filamentous particles. For example, by interfering with filamentous-particle assembly, stability, or internalization into cells, we would be eliminating this viral persistence strategy (Figure S1B and 7C). Combining an approach that targets filamentous virus particles with membrane-fusion inhibitors, therapeutic antibodies, or vaccination might prolong the effectiveness of treatments or immunizations focused on the viral cell-entry machinery. For example, several monoclonal antibodies targeting the ectodomain of the influenza proton channel M2 (M2e) have been shown to prevent filamentous particle assembly and to fragment filamentous virus particles (Kolpe et al., 2019; Roberts et al., 1998; Rossman et al., 2010). M2 is a minor component of the virion surface, and M2e-targeting antibodies arise infrequently during a natural course of infection or in response to vaccination. Specific stimulation of the immune system to enrich for both M2e- and HA-targeting broadly neutralizing antibodies, might thus present a viable strategy to achieve lasting immunity. In another example, NA inhibitors were shown to disproportionately inhibit the release of filamentous influenza particles during virus budding from infected cells because of the higher relative NA-content of spherical particles (Vahey and Fletcher, 2019). Combining HA and NA inhibitors might thus delay resistance in an unanticipated way because NA pressure increases the proportion of spherical particles, ready targets for HA inhibitors. By exposing a concealed viral strategy of adaptation, this work also exposes a previously unknown viral weakness enabling novel antiviral strategies.

## ACKNOWLEDGEMENTS

We thank Goran Bajic (Harvard Medical School) for help with Fab expression and purification, Milos Popovic (Boston University) for help with analysis codes, and Stephen C. Harrison (Harvard Medical School), Bruce L. Goode (Brandeis University), Priscilla L. Yang (Stanford University), and Amy S.Y. Lee (Brandeis University) for comments on the manuscript. We acknowledge support from the NIH Director’s New Innovator Award 1DP2GM128204 (to TI), the NSF MRSEC DMR-1420382 (to TI), and the NIH grant R01AI134824 (to KC).

## AUTHOR CONTRIBUTIONS

TL, Influenza virus and EBOV: Experiment design, Acquisition of data, Analysis and interpretation of data; ZL, ED, ML, Influenza virus: Acquisition of data, Analysis and interpretation of data; EM, KC, EBOV: VLP design, endosomal fusion-experiment design, construct generation; TI, Conception and design, Experiment design, Acquisition of data, Analysis and interpretation of data; TI, TL, Writing of the manuscript. All authors have contributed to manuscript revision.

## STAR METHODS

### Reagents

#### Cells

MDCK.2 cells (ATCC strain CCL-34) and Hela cells were propagated in DMEM supplemented with 10% FBS. All infections were performed in OptiMEM (ThermoFisher Scientific) with or without 1 µg/ml TPCK-trypsin (Sigma) or M-Fab. Human embryonic kidney 293F cells were a generous gift from Stephen C. Harrison, Harvard Medical School. 293F cells were maintained in FreeStyle 293 Expression Medium (ThermoFisher Scientific), and used for the expression of M-Fab, HC19 IgG, KZ52 IgG and Ebola VLPs.

#### Antibodies

Hybridomas for SiteA and A2 antibodies were a generous gift from Judith M. White, University of Virginia. Purified LC89 antibody was a generous gift from Stephen A. Wharton, MRC National Institute for Research, London, UK. SiteA antibody binds an epitope on the HA1 head accessible on both the pre- and post-fusion HA. A2 detects an epitope on the low-pH HA1, and LC89 detects the low-pH helix-to-loop transition in HA2, residues 106-112 (Wharton et al., 1995); both are reactive against HA0 and either HA1 or HA2 in Western blot.

### Influenza viruses

Influenza viruses have A/Aichi/68 (X31) HA and the remainder of the segments from A/Udorn/72 (Ivanovic et al., 2012). D112_2_A mutant was characterized previously (Ivanovic et al., 2013). H17_1_Y mutant was made and grown according to our D112_2_A protocol (Ivanovic et al., 2013), except that it was passaged at a higher multiplicity of infection (MOI) of 0.1 before the final, MOI 12 infection that was not modified. Uncleaved virus was generated as described for the cleaved virus (Ivanovic et al., 2012) except that trypsin was omitted in the final, MOI 12 infection before purification. Purifications of the short WT, D112_2_A, and H17_1_Y viruses used in Figure 5A and B were done according to our previous short-particle purification protocol on 20-60% sucrose gradient (Ivanovic et al., 2012), and the size-distributions of the resultant fractions were indistinguishable from the SUP fractions developed here. For the pelleting protocol, viruses were passed through a 30% sucrose cushion as before (Ivanovic et al., 2012), and then subjected to 8 cycles of centrifugation at 3250 rcf for 1.5 hrs at 4 °C. For each spin, virus was diluted to 900 µl HNE20 (20 mM HEPES NaOH pH 7.4, 150 mM NaCl and 0.2 mM EDTA). After each spin, 850 µl of the supernatant volume was removed, and the pellet was resuspended in HNE20 over 6-14 hours at 4 °C. The final, 8^th^-spin pellet constitutes the PELLET fraction. Supernatants were combined and concentrated by a 1-hr spin at 100,000 xg (the SUP fraction).

All viruses were stored in HNE20. All purified viruses were sequenced to confirm HA-segment correctness. Completeness or near-absence of HA cleavage/activation on the purified cleaved or uncleaved virus preps was verified by A2-antibody Western blots (Ivanovic and Harrison, 2015). Each virus passage and each virus manipulation (e.g. fluorescent labeling, pelleting fractionation) were followed by measurements of specific virus infectivity using a combination of the standard plaque and hemagglutination (HA) assays to ensure a constant ratio of plaque forming units (PFU) to HA units (HAU reports on the physical virus-particle count independently of the particle morphology – see under Electron microscopy). A typical PFU/HAU ratio for the viruses is ∼1-5E4. Total protein concentration of the purified virus preps was determined by the Bradford assay (BioRad). For hemagglutination inhibition assay (HAI), viruses were preincubated with HC19 IgG at room temperature for 30min, then the standard HA assay was performed in the presence of HC19 IgG.

### Ebola VLPs

The cDNA sequence of EBOV GPdelMuc (EBOV ‘Mayinga’, GenBank accession number AF086833, lacking the mucin-like domain including the amino acids 309-489 (Jeffers et al., 2002)) was expressed under a CMV promoter by insertion into the mammalian expression vector pCAGGS. A cDNA encoding the EBOV matrix protein VP40 was similarly cloned into the expression vector pCAGGS. EBOV VP40 N-terminally linked to a Renilla luciferase was generated as described previously (Martin-Serrano et al., 2004) by replacing the eGFP coding sequence with a luciferase coding counterpart followed by cloning into pCAGGS.

For the EBO VLP expression, 1E7 293F cells were co-transfected with 4.5 µg of pCAGGS-Renilla luciferase-VP40, 0.5 µg of pCAGGS-VP40, and 5 µg of pCAGGS-GPdelMuc using 15 µg of polyethylenimine (PEI). Culture supernatant was harvested 48 hours post-transfection and clarified by centrifugation at 2000xg for 10min. VLPs were concentrated by ultracentrifugation at 154,000xg for 3 hours, then pelleted through a 20% sucrose cushion in HNE20 at 230,000xg for 2 hrs. The VLP pelleting protocol to generate EBO VLP PELLET and SUP fractions was similar to that used for influenza viruses, except that the spins were done at 2142 xg.

### Electron microscopy

Influenza virus PELLET fractions were adjusted to 2e4 HAU/ml, and SUPs and short gradient-purified fractions were adjusted to 1e5 HAU/ml. As a precaution, viruses were pretreated with 25 µM rimantadine (M2 inhibitor, Sigma) for 30 minutes at room temperature (RT) to prevent fragmentation of the filamentous particles (Rossman et al., 2012). 2% phosphotungstic-acid staining was performed as previously described (Ivanovic et al., 2012). Images were taken on Philips Morgagni Version3.0 TEM (80 kV, 1k CCD) using AMT Camera Version 600.335o software. Particle-size measurements were performed using custom MATLAB codes as previously described (Ivanovic et al., 2012).

To ensure that HA-assay measurements accurately report on virus-particle counts for the influenza PELLET and SUP fractions, we performed EM-based particle counting as follows. PELLET and SUP fractions diluted as above were each mixed with a fixed concentration of purified reovirus particles. Reovirus particles have regular icosahedral structures ∼85-nm in diameter (Zhang et al., 2005), and appear as distinct, regular, darker objects, clearly distinguishable from influenza virus particles in the negative-stain electron micrographs. 1000-3000 total particles were counted to determine the SUP/reo and PELLET/reo particle ratios. The resultant value for PELLET/SUP particle ratio closely matched (within 30%) the PELLET/SUP particle ratio derived from their respective input HAU/ml. We therefore used HA-assay values to represent a measure of input physical particles in experiments independent of the particle morphology.

Electron Microscopy of the EBO VLP SUP and PELLET were performed similarly, with input particle numbers adjusted based on the images seen under the microscope. A fixed concentration of reovirus particles were mixed with EBO VLP fractions to determine the relative particle concentration between the SUP and PELLET fractions. Same number of particles for the EBO VLP PELLET and SUP were used in endosomal-fusion experiments (Figure 6C).

### IgG expression and purification

The expression vectors (modified pVRC8400) for HC19 light and heavy chains were a generous gift from Stephen C. Harrison, Harvard Medical School. The expression vectors (pMAZ) for KZ52 light and heavy chains were previously described (Maruyama et al., 1999). IgGs were produced by transient transfection of 293F cells with PEI (0.4 µg heavy chain DNA: 0.6 µg light chain DNA: 1.5 µg PEI: 1E6 cells) and purified from the cell-culture supernatant 7 days post transfection using protein G resin (GE Healthcare).

### M-Fab expression, purification and labeling

The expression vectors (modified pVRC8400) for MEDI8852 Fab light and heavy chains were a generous gift from Stephen C. Harrison, Harvard Medical School. The heavy-chain construct contains the C-terminal 6xHis tag. For the fluorescent labeling of M-Fab, we modified the heavy-chain construct to also include a Sortase recognition motif LPETGG between the heavy-chain sequences and the 6xHis tag. Fab fragments were produced by transient transfection of 293F cells with PEI (1 µg DNA: 1.5 µg PEI: 1E6 cells). Fabs were purified from the culture supernatants 5 days post transfection by a passage over a HisTrapFF column (GE Healthcare) followed by gel filtration chromatography on Superdex 200 Increase (GE Healthcare) in PBS.

A 10-fold molar excess of GGGK peptide (GenScript) was labeled with JF549-NHS-ester dye (Tocris Bioscience) in 100 mM sodium bicarbonate (pH 8.4) buffer. The desired peptide, singly labeled at the lysine residue was purified by reverse-phase HPLC on a C18 column (Grace) and its identity was confirmed by LC/MS and by its reactivity with LPETGG-M-Fab in the Sortase-labeling reaction. 6xHis-tagged SortaseA5 was expressed in *E.coli* BL21(DE3) cells and purified as previously described (Chen et al., 2011). A 1: 1: 25 molar ratio of SortaseA5: LPETGG-M-Fab: GGGK-JF549 was used for the Sortase-labeling reaction in PBS supplemented with 5 mM CaCl_2_ and 5 mM MgCl_2_ for 30 minutes at RT. Unreacted LPETGG-M-Fab and Sortase were removed from the labeled product using NiNTA resin in PBS containing 10-30 mM imidazole buffer. M-Fab-JF549 product was buffer exchanged into PBS.

### Stochastic simulations

We used our previously published simulation codes without modification(Ivanovic and Harrison, 2015). Simulation were run using the following input parameters: 1) *k*_*e*_ = 0.003sec^-1^, where *k*_*e*_ is the rate of fusion peptide release, 2) *f*_un_ = 0.5, where *f*_un_ is the unproductive HA fraction, 3) N_h_ = 3, where N_h_ is the number of HA neighbors in the fusion cluster, 3) *f*_in_ = 0 to 0.95, where *f*_in_ is the inactive HA fraction in 0.05-0.1-value increments, and 4) input values for patch size of 50, 100, 200, 500, 1000, 3000, or 30000, to obtain actual patch sizes of 55, 121, 235, 583, 1159, 3463, or 34627, respectively. The shape of the contact patch in the simulation was kept as a circle because both unproductive and inactive HAs subdivide the area into small segments rendering the overall starting shape irrelevant for the measured outcomes.

### Single-particle experiments

#### Hemifusion

We used the protocol for influenza membrane fusion described previously(Ivanovic et al., 2013) with modifications. Virions were labeled in 25-µl reactions at 0.24 mg/mL with 1,1’-Dioctadecyl-3,3,3’,3’-Tetramethylindodicarbocyanine, 4-Chlorobenzenesulfonate Salt (DiD, ThermoFisher Scientific) at 10 µM for 1.5 hours at RT. Labeled virions were used within one week. Liposomes used for planar bilayer generation consisted of 4:4:2:0.1:2 × 10^−4^ ratio of 1, 2, dioleoyl-sn-glycero-3-phosphocholine (DOPC) (Avanti Polar Lipids), 1-oleoyl-2-palmitoyl-sn-glycero-3-phosphocholine (POPC; Avanti Polar Lipids), sn-(1-oleoyl-2-hydroxy)-glycerol-3-phospho-sn-3’-(1’-oleoyl-2’-hydroxy)-glycerol (ammonium salt) (18:1 BMP (R,R)) (Avanti Polar Lipids), bovine brain disialoganglioside GD1a (Sigma), and N-((6-(biotinoyl)amino)hexanoyl)-1,2-dihexadecanoyl-sn-glycero-3-phosphoethanolamine (biotin-X DHPE) (Molecular Probes, Life Technologies), where sialic acid residues on GD1a served as receptors for influenza virions. Briefly, as before(Ivanovic et al., 2013), planar bilayers were generated from 200-nm liposomes by vesicle-spreading method within the channels of a PDMS flow cell. They were labeled with fluorescein-conjugated streptavidin (30 µg/ml, Invitrogen) that served as an external pH-probe. DiD-labeled virions (untreated or treated with M-Fab for 20 min at RT) were attached for 5-15 min. Unattached virions were washed with HNE20, and the pH was dropped with pH-4.8 or pH-5.2 buffers (10 mM citrate, 140 mM NaCl and 0.1 mM EDTA). For M-Fab-treated samples, M-Fab was included in the low pH buffer, and after 3 min of the low-pH buffer flow at 60 µl/min, the flow was reduced to 6 µl/min to preserve M-Fab. We followed hemifusion as a spike in individual virion fluorescence resulting from DiD-fluorescence dequenching upon dilution in the target membrane. All experiments were performed at 23±0.4 °C.

#### Fluorescent M-Fab binding

Cleaned and plasma etched slides (Ivanovic et al., 2013) were silanized with N-(2-aminoethyl)-3-aminopropyltrimethoxysilane, amino silane (United Chemical Technologies) and then PEGylated with a mixture of biotin-PEG-SVA-5000 and mPEG-SVA-5000 (1:40 molar ratio; Laysan, Inc.) according to a published protocol (Diao et al., 2012). They were dried and stored protected from light and moisture at −80 °C until use. For fluorescent M-Fab binding experiments, PDMS flow cells were assembled as before(Ivanovic et al., 2013) and the following sequence of steps was followed each time: 1) PBS wash, 5 min at 30 µl/min; 2) 5-min incubation with 0.2 mg/ml NeutrAvidin (ThermoFisher Scientific); 3) PBS wash, 2 min at 30 µl/min; 4) 6-min incubation with biotinylated SiteA antibody (0.1 mg/ml); 5) PBS then HNE washes, 2 min at 30 µl/min; 6) imaging for background subtraction; 7) 7-min incubation with unlabeled virions pre-incubated with M-Fab-JF549 for 20 min at RT; 8) HNE20 wash, 30 sec at 30 µl/min; 9) imaging for intensity measurements, pH7.4 (4 fields of view); 10) pH-4.8 buffer wash, 3-min at 60 µl/min, and incubation for the total of 20 min; 11) imaging for intensity measurements, pH4.8 (4 fields of view, including one shared with pH7.4-imaging). When SiteA antibody was replaced by a non-specific mouse antibody, a very small amount of nonspecific sticking of influenza virus particles was observed (∼1%).

#### Microscope configuration

The excitation pathway was unchanged relative to our previous setup(Ivanovic et al., 2013), except that we now added a 647-nm laser (Obis, Coherent) and the diameter of the illuminated area was approximately 136 µm. Total internal reflection (TIR) illumination was achieved using a commercial micro-mirror platform, RM21 (Mad City Labs, Inc.) essentially as described (Larson et al., 2014). Briefly, the system spatially separates the excitation and emission light within the objective lens and eliminates the need for a dichroic. An integrated XY micro-positioning, and XYZ nano-positioning stage is designed around a fixed objective lens (Olympus 60X ApoN NA=1.49) with the open access to its back aperture required for micro-mirror-based illumination. The system was additionally equipped with a 180-mm achromatic tube-lens. Image was projected onto an entire 512 × 512 pixel EM-CCD sensor (Model C9100-13; Hamamatsu) for the final pixel representation of 160 nm.

For the single-particle hemifusion experiments, 488-nm laser (Obis, Coherent) was used to excite fluorescein, and 647-nm laser (Obis, Coherent) was used to simultaneously excite DiD. The power of the 488-nm laser was 1-2 µW, and the 647 laser was 0.25-1 µW over the illuminated area. Exposure time was between 0.4 and 0.8 sec, depending on the virus and the inhibitor concentration. For the fluorescent intensity measurements of M-Fab-JF549 binding to virus particles, the power of the 552-nm laser was 5.1 µW over the illuminated area, and the exposure time was 1 sec.

### Data analysis

All data analysis was done in MATLAB (MathWorks).

#### Single-particle hemifusion

Particle centroids, pH-drop time, and DiD-dequenching times for individual particles were extracted from time-lapse movies as described (Floyd et al., 2008). We defined hemifusion yield as the percent of detected particles that hemifused within 20 minutes of pH drop. Probability-density distributions of hemifusion lag times were fitted with gamma distribution to derive mean lag-times (Floyd et al., 2008; Ivanovic and Harrison, 2015; Ivanovic et al., 2012; Otterstrom et al., 2014).

#### Derivation of long-particle hemifusion kinetics

Hemifusion-yield curves for PELLET and SUP particles (Figure 3D) were scaled to adjust the no-M-Fab hemifusion to 100%. We justify this in the context of our previous analysis, which found that the lower than 100% yield in the absence of HA inhibition reports on the efficiency of the assay, but the probability of fusion-cluster formation is 100% (Ivanovic and Harrison, 2015). This assumption is further justified by PELLET maintaining unreduced hemifusion yield at 67 nM M-Fab; if fusion-cluster formation was already limiting for no M-Fab, HA inactivation would cause a further reduction in hemifusion yield (Figure 4B) (Ivanovic and Harrison, 2015). In what follows, we illustrate with one example how the scaled yield-data were then processed to derive long-particle yields. At 268 nM M-Fab, there is 80% (Figure 3D) and 70% (not shown, scaled data from Figure 3C) reduction in hemifusion yield for SUP and PELLET, respectively. Since SUP-like particles represent approximately 50% of the PELLET (Figure 1B), the loss of fusing particles determined for the SUP can account for 40% loss in the PELLET; the remaining 30% must have resulted from inhibition of long particles, and the long-particle yield is thus 70% (Figure 3D). For samples where PELLET yield-reduction was not greater than that predicted from the SUP data, long-particle yield was set to 100%.

We performed bi-gamma fitting of the PELLET lag-time distributions (Figure 3B, *bottom*) using the following formula,

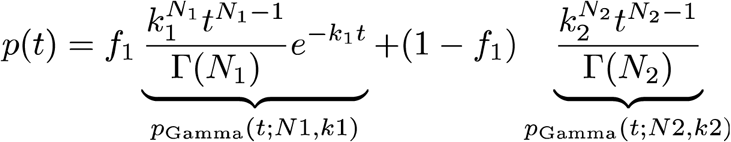

Here, *p*_Gamma_(t; N,*k*) is the probability-density function of the Gamma distribution with rate *k* and shape parameter N, and *f*1 is the fraction of population whose distribution is represented by (N1,*k*1) or the SUP-like particles, with the remainder sampling a (N2,*k*2) or the long-particle distribution. *f*1, *k*1, and N1 values were fixed, and N2, and *k*2 were derived from the fits. *k*1 and N1 were derived from single-gamma fitting of the SUP data (Figure 3B, *top, red lines*). *f*1was calculated as follows. From the SUP- and long-particle yield values (Y_SUP_ and Y_PELLET_) (Figure 3C), we calculated the fraction of the observed hemifusion events contributed by the long particles in the PELLET, *f*_long_ = Y_long_ / (Y_long_ * 0.5 + Y_SUP_ * 0.5). The two yield values were multiplied by 0.5 because SUP-like particles represent approximately 50% of the PELLET (Figure 1B). *f*1 was fixed at (1-*f*_long_). We show the bi-gamma fit curves (*black lines*), and the underlying curves for long (*blue lines*) and SUP-like particles (*red lines*) in the PELLET (Figure 3B, *bottom*).

#### Particle fluorescence-intensity measurements

The intensity-analysis codes are supplied: 1) s_intensity_analysis_of_imageCB.m, 2) s_intensity_analysis_of_image_step2.m, and 3) s_intensity_analysis_of_image_step3.m. The first code averages the first 10 frames of the particle movies (M-Fab-JF549 particles) and the no-particle movie (background), and subtracts the background image from the averaged particle images. The second code plots the background-processed image and prompts the user to manually draw regions of interest (masks) around particles whose intensity is to be quantified. Manual masks were preferred in this case because of the long particles that can be mistaken for multiple particles by automated particle-detection codes. The third code then integrates intensities within the masks.

### Infectivity measurements

#### scTCID50

Virus was pre-incubated with or without M-Fab in OptiMEM for 30 minutes at RT, then serially diluted 4-fold in the same media. Trypsin was omitted to prevent activation of the released virus and new rounds of infection. Confluent monolayers of MDCK.2 cells in 96-well plates were washed with HBSS twice, and 100 µl of the virus dilutions were added to wells. The highest input virus concentration was 2-7*10^3^ HAU/ml. For scTCID50 that included EIPA, cells were pretreated with 50µM EIPA for 1 hour and EIPA was included in the media for the duration of infection. Cells were fixed 24 hrs post infection with 3.7% formaldehyde, then stained with crystal violet. Plates were scanned using Gel Doc (Biorad), and the extent of cell death in each well evaluated by densitometry using ImageJ. The densitometry value in wells with complete cell death (background) was subtracted from all measurements, and the amount of cell death quantified relative to wells with no cell death. scTCID_50_ titer is derived from the resultant plots of cell death versus particle concentration as the inverse of the particle concentration that resulted in 50% cell death. Relative reduction in infectivity was quantified as the ratio of scTCID_50_ values for the untreated (baseline) and M-Fab-treated samples. We obtained the same baseline titers (tested with WT and H17_1_Y PELLET and SUP) when new rounds of infection were additionally prevented by the addition of 20 mM ammonium chloride(Daniels et al., 1985) 4-hours post infection, confirming that our experimental setup resulted in a single-cycle of infection. For each experiment, each condition was done in duplicate. At least three independent experiments were performed. The same results were obtained with a second clone of each WT and mutant viruses.

#### Modified plaque assay

Uncleaved PELLET and SUP viruses were diluted to 500 HAU/ml (for the uncleaved/M-Fab treated samples) or 50 HAU/ml (for the remaining samples) in OptiMEM for pre-treatment. For cleaved samples, uncleaved viruses were pre-treated with 1 µg/ml TPCK-trypsin (Sigma) for 1 hr at RT, and trypsin was inactivated with 20 µg/ml soybean trypsin inhibitor (Sigma). For M-Fab pre-treatment, precleaved or uncleaved viruses were incubated with 268 nM M-Fab for 30 min at RT. The pre-treatment reactions were directly used as the first virus dilution in the subsequent plaque assay, and were serially diluted 10-fold in OptiMEM (no additives) for 5 more steps. 100 µl of the virus dilutions were attached to confluent monolayers of MDCK.2 cells in 6-well plates for 1 hour at RT. Unattached virus and free M-Fab were washed twice with HBSS, and viruses were allowed to internalize for 4 hours at 37 °C in OptiMEM (no additives). Media was then replaced with OptiMEM/0.6% Oxoid Agar/1 µg/ml TPCK-trypsin, and the plates were incubated for 2.5 more days at 34 °C for plaque formation. Removal of M-Fab after pre-incubation and virus attachment, and addition of trypsin only after virus internalization, ensure that any differences among samples are due to differences in the initial infection by SUP and PELLET. An additional control had trypsin present in all virus dilutions and during the 4-hour internalization phase. No infectivity difference was detected between this sample and the cleaved sample where trypsin was inactivated after digestion, confirming complete virus activation by the 1-hour trypsin treatment. The cleaved-virus reference in the Western blot (Figure 4C) was a fully cleaved virus prepared in a standard way (not deriving from the uncleaved virus prep). The Western blot used LC89 antibody for HA0 and HA2 detection.

### Estimate for the fraction of HA inactivation by M-Fab and for particles with uncleaved HA

#### Particles with uncleaved HA

To estimate the extent of HA cleavage on purified ‘uncleaved’ virus particles we compared the HA2 band intensity in immunoblot for a range of input total-protein amounts, to the HA2 band intensity of two input total-protein amounts for purified, fully cleaved particles (Figure 4C). Both input total-protein amounts were determined by Bradford assay (BioRad). We estimate that less than 1% of HA is cleaved on ‘uncleaved’ virus particles.

#### M-Fab-bound particles

We estimate that about 75% of HAs are inactivated at 268 nM M-Fab. This value derives from the comparison of the single-particle hemifusion-yield data for the SUP and long particles in the PELLET (Figure 3D) to the fusion-cluster yield results from stochastic simulations for the PS 121 and PS 583 (Figure 4B, orange and blue lines). We estimate that the average particle in the SUP has a patch size containing 121 HAs (Figure 1B, insets) (Ivanovic et al., 2013) and the long particles in the PELLET are ∼5-times longer (Figure 1B, insets) corresponding to ∼5-times larger contact patch (corresponding to PS 583). At 268 nM M-Fab, scaled hemifusion yield for the SUP is ∼20%, and for the PELLET about ∼70% (Figure 3D). These values correspond to ∼75% HA inactivation, or only ∼25% active HAs (Figure 4B, orange and blue lines).

#### Particles with uncleaved HA during endosomal entry

We estimate that less than 25% of HAs get processed during endosomal entry by uncleaved particles because they are less infectious than cleaved particles at 268 nM M-Fab (Figure 4D) for which we estimated that ∼25% HAs are active (see *M-Fab-bound particles*). Taking 20% as an estimate for active HAs during endosomal entry by uncleaved particles in the absence of M-Fab, we estimate that there are only ∼5% (0.25*0.2=0.05) active HAs remaining during endosomal entry of uncleaved particles in the presence of 268 nM M-Fab.

### Ebola VLP luciferase assay

Hela cells were seeded on polystyrene 96-well plates (Fisher Scientific) 24 hours before the experiment. The same number of particles for EBO VLP SUP and PELLET were preincubated in the absence or presence of KZ52 IgG for 30 min before attachment to cells. Attachment was done by spinoculation at 2000 xg for 2 hours at 4°C. Cell were then incubated at 37°C for 5 hours in the presence of KZ52. At this point, EnduRen live cell luciferase substrate was added to cells according to manufacturer’s protocol (Promega), and the cells were returned to 37°C incubator for an additional 1.5 hours before the detection of the luminescence with Promega GloMax luminometer (Promega). Each experiment was performed in duplicate, and 3 independent experiments were performed. The uninhibited SUP and PELLET signals were ∼30-fold and ∼15-fold over the cell-only negative control, respectively, and the data in Figure 6 report the percent signal for treated samples relative to the untreated samples.

## SUPPLEMENTAL INFORMATION

**Figure S1.**
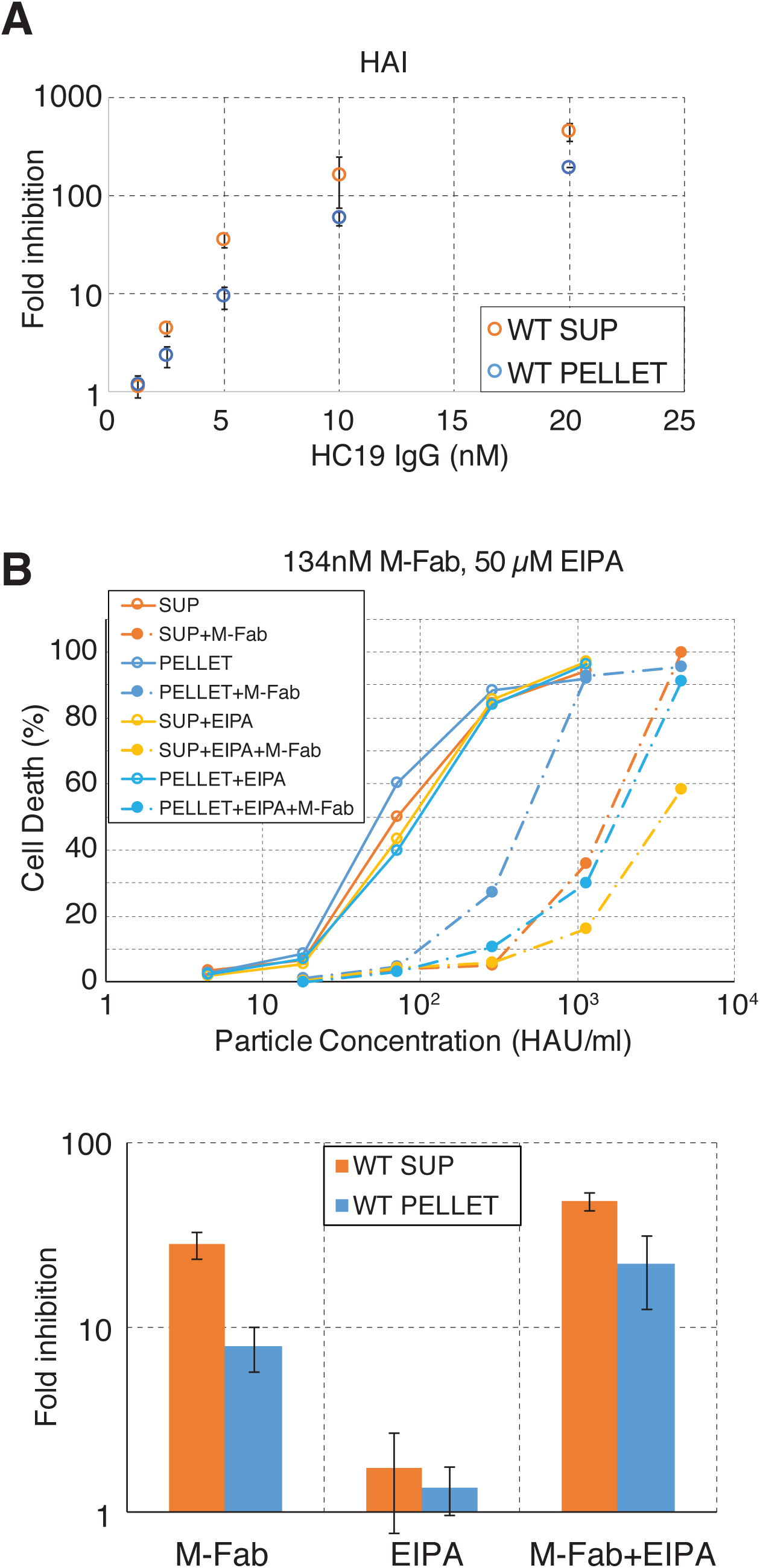
Control experiments explaining the functional advantage of the PELLET in the presence of HC19 IgG or M-Fab. (A) PELLET particles are less sensitive than SUP particles to inhibition of cell-attachment by HC19 IgG. Hemagglutination inhibition (HAI) was calculated relative to a no-HC19 IgG control. (B) Long particles in the PELLET are responsible for the functional advantage of the PELLET in the presence of M-Fab. A sample scTCID_50_ experiment (*top*), and the average and standard deviation from three independent scTCID_50_ experiments (*bottom*) are shown. An inhibitor of macropinocytosis EIPA eliminates the functional advantage of the PELLET in the presence of M-Fab.

**Figure S2.**
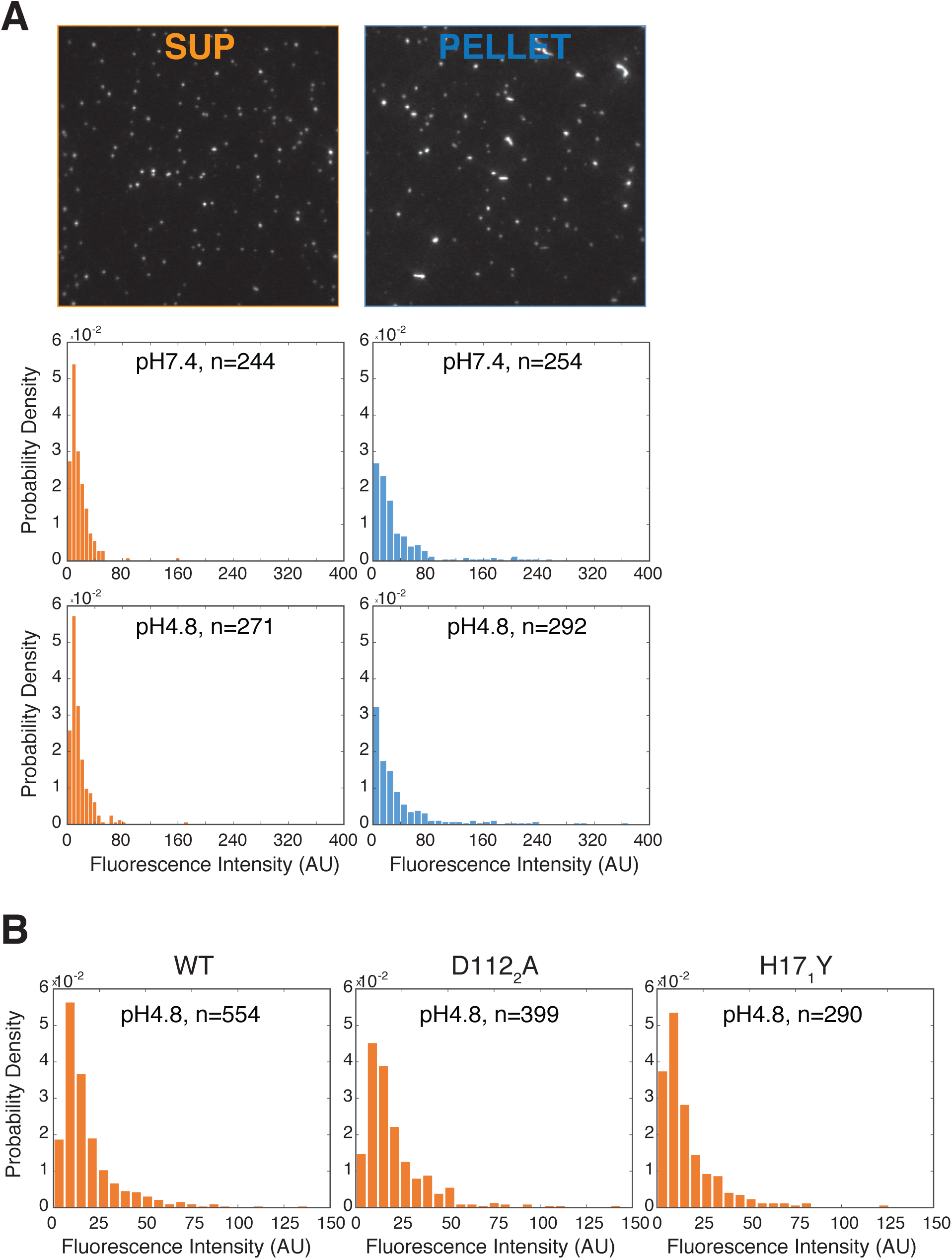
Quantification of M-Fab^fl^ binding to unlabeled particles. Glass coverslips were passivated with a mixture of polyethylene glycol (PEG) and PEG-biotin. Biotinylated antibody targeting HA1 is specifically captured with neutravidin, and it recruits virus particles for imaging. Virus particles were pre-incubated with 134 nM M-Fab^fl^ before capture, and unbound M-Fab^fl^ was washed away after capture. (A) Representative particle-images (*top*), and particle fluorescence-intensity quantification at neutral pH (*middle*), or after 20 minutes at pH 4.8 (*bottom*). Data were pooled from several experiments all acquired on the same day. The number of particles represented by the histograms (n) is indicated on the plots. (B) Fluorescence-intensity quantification for short WT, D112_2_A, and H17_1_Y particles after 20 minutes at pH 4.8.

**Figure S3.**
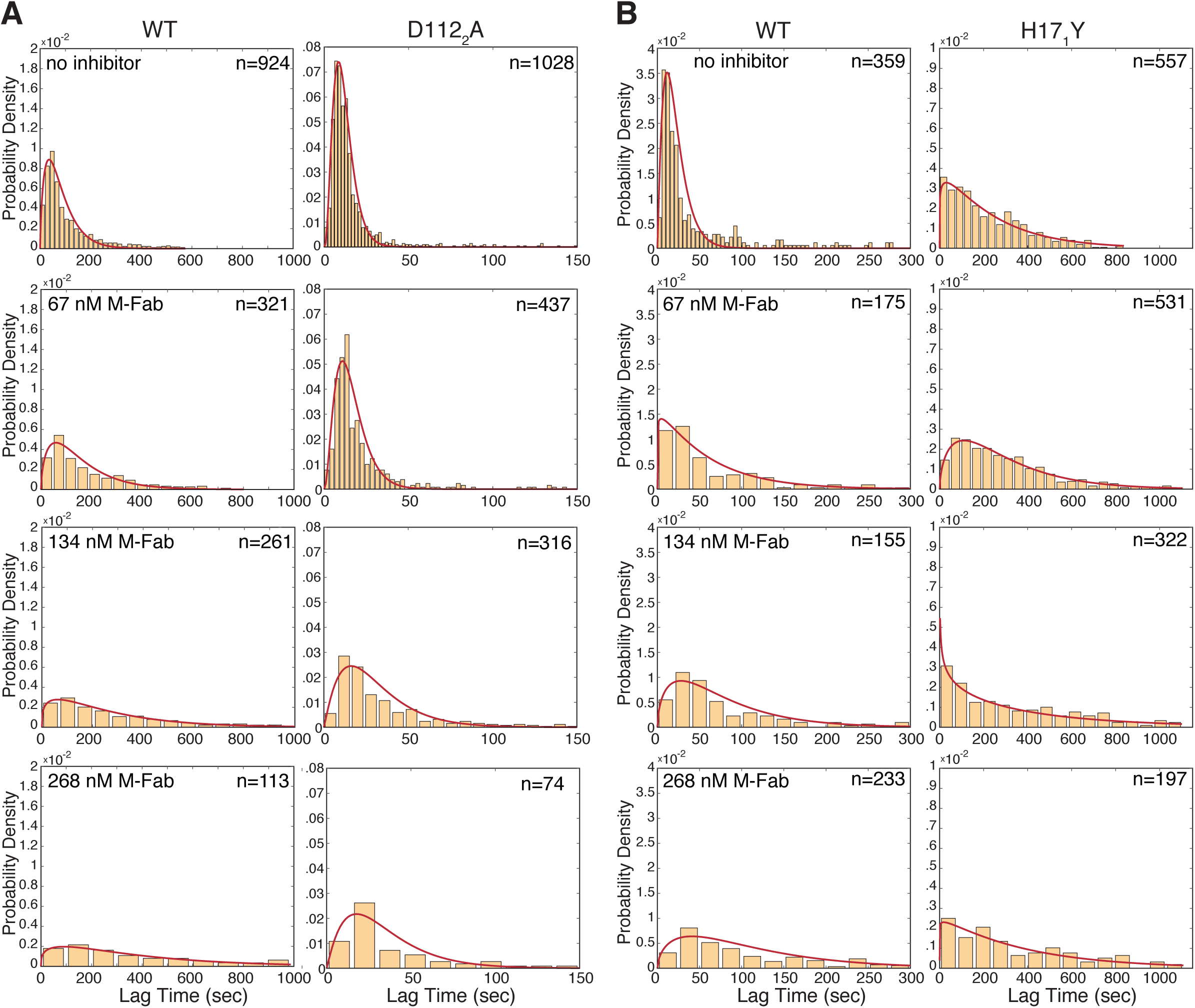
Single-particle measurements of membrane fusion for short D112_2_A and H17_1_Y particles. Hemifusion lag times with gamma-distribution fit (*red line*). Data from several experiments were pooled where each condition was represented on each day. The total number of particles (n) represented by each histogram is indicated. (A) Short WT and D112_2_A particles, pH 5.2 (B) Short WT and H17_1_Y particles, pH 4.8.

**Figure S4.**
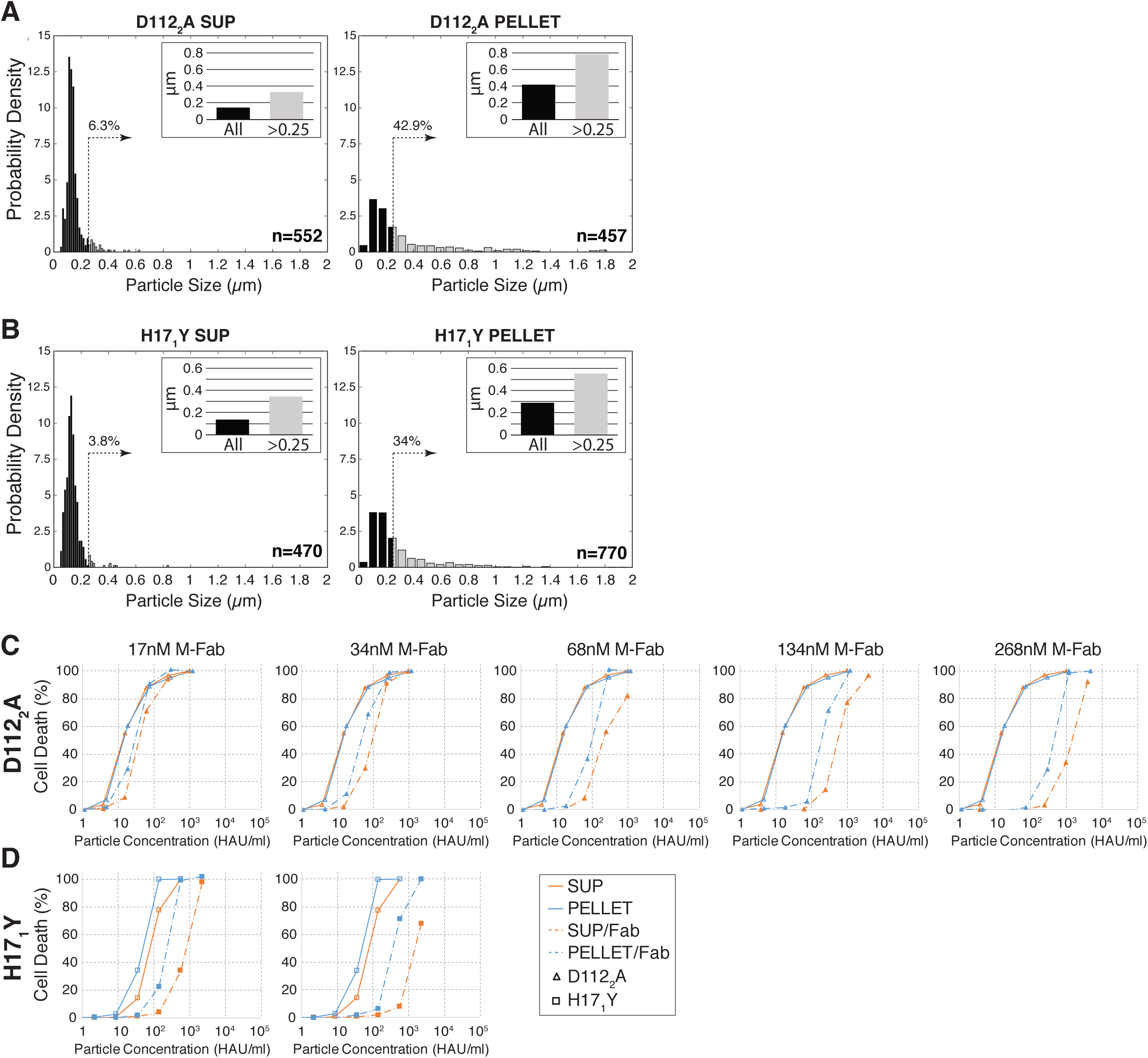
D112_2_A and H17_1_Y particle fractionation and scTCID_50_ measurements. (A and B) Particle-length distributions derived from electron micrographs like those shown in Figure 1A. Particle-length data were normalized so that the total area under the bars is equal to 1 on all plots. Dotted vertical lines mark 0.25 µm length cutoff, and the percentage of particles longer than 0.25 µm is indicated above. Each inset shows mean particle length for the entire population (black bar), or the subset of the population longer than 0.25 µm (gray bar). (C and D) Sample D112_2_A (C) and H17_1_Y (D) cell-death data plotted in Figure 5C-E. The no-inhibitor baseline values, shown on all plots, were used in Figure 5C, or as values from which to calculate infectivity shifts in the presence of M-Fab in Figure 5D-E.

**Movie S1 | A sample movie of SUP hemifusion at pH 5.2 and no M-Fab.** Exposure time is 0.5 sec and the movie is at 80 frames per second (fps).

**Movie S2 | A sample movie of the PELLET hemifusion at pH 5.2 and no M-Fab.** Exposure time is 0.5 sec and the movie is at 80 fps.

**Movie S3 | A sample movie of the SUP hemifusion at pH 5.2 and 134 nM M-Fab.** Exposure time is 0.8 sec and the movie is at 50 fps.

**Movie S4 | A sample movie of the PELLET hemifusion at pH 5.2 and 134 nM M-Fab.**

Exposure time is 0.8 sec and the movie is at 50 fps.

**Code S1.** s_intensity_analysis_of_imageCB.m (see Materials and Methods).

**Code S2.** s_intensity_analysis_of_image_step2.m (see Materials and Methods).

**Code S3.** s_intensity_analysis_of_image_step3.m (see Materials and Methods).

